# Functional shifts of soil microbial communities associated with *Alliaria petiolata* invasion

**DOI:** 10.1101/2020.08.13.248849

**Authors:** Katherine Duchesneau, Anneke Golemiec, Robert I. Colautti, Pedro M. Antunes

**Affiliations:** Department of Biology, Queen’s University, Kingston, Ontario, K7L 3N6; Department of Biology, Algoma University, Sault Ste-Marie, Ontario, P6A 2G4

**Keywords:** *garlic mustard*, microbial function, plant–soil (below-ground) interactions, arbuscular mycorrhiza, invasion ecology, mechanisms of plant invasion

## Abstract

Soil feedback is thought to be an important contributor to the success of invasive plants. Despite evidence that invasive plants change soil microbial diversity, the functional roles of microbes impacted by invasion are still unclear. This knowledge is a critical component of our understanding of ecological mechanisms of plant invasion. Mounting evidence suggests *Alliaria petiolata* can suppress arbuscular mycorrhizal fungi (AMF) to disrupt native plant communities in controlled laboratory and greenhouse experiments, though it is less clear if allelochemicals persist under natural field conditions. Alternatively, invasive plants may accumulate pathogens that are more harmful to competitors as predicted by the Enemy of my Enemy Hypothesis (EEH). We examined changes in functional groups of soil bacteria and fungi associated with ten naturally occurring populations of *A. petiolata* using amplicon sequences (16S and ITS rRNA). To relate soil microbial communities to impacts on co-occurring plants, we measured root infections and AMF colonization. We found no changes in the diversity and abundance of AMF in plants co-occurring with *A. petiolata*, suggesting that mycorrhizal suppression in the field may not be as critical to the invasion of *A. petiolata* as implied by more controlled experiments. Instead, we found changes in pathogen community composition and marginal evidence of increase in root lesions of plants growing with *A. petiolata*, lending support to the EEH. In addition to these impacts on plant health, changes in ectomycorrhiza, and other nutrient cycling microbes may be important forces underlying the invasion of *A. petiolata* and its impact on ecosystem function.

## Introduction

Evolutionary history constantly reshapes the dynamic interactions between micro- and macro-organisms, determining the composition of plant terrestrial communities over time (Bevers et al., 2015; Reynolds, Packer, Bever & Clay, 2003; van Der Heijden, Bardgett & van Straalen, 2008; Wagg, Bender, Widmer & van der Heijden, 2014). Human activities facilitate the global movement of species that can invade and disrupt these complex ecological networks, often with unforeseen impacts on native species diversity and ecosystem processes. For example, the Asian plant *Pueraria montana* (kudzu), introduced to control erosion in the southern United States, affects neighboring plants directly as a structural parasite and indirectly by increasing soil nitrogen availability, thereby excluding plants adapted to low soil nitrogen conditions (Forseth & Innis, 2004). Consequently, it is important to understand the wide range of ecological impacts caused by invasive plants in order to predict ecosystem-level responses.

The disruption of soil microbial communities is thought to be a major contributor to the expansion of invasive plants (Callaway, Thelen, Rodriguez & Holben, 2004; Klironomos, 2002; Wolfe & Klironomos, 2005; Waller et al., 2020; Zobel & Öpik, 2014). Although not all invasive species are drivers of ecosystem changes (see Bauer, 2012; MacDougall & Turkington 2005), some can selectively disrupt microbes to gain an advantage over their native competitors. For example, two hypotheses on how invasive plants can modify the soil microbiome to outperform co-occurring native plants are linked to the disruption of native mutualisms (Stinson et al., 2006) and accumulation of native pathogens (Flory & Clay, 2013; Stricker, Harmon, Goss, Clay & Flory, 2016). Thus, to determine how shifts in microbial communities can change ecosystems, it is important to recognize the different functional roles of distinct microbial taxa (Nguyen et al., 2016; Petchey & Gaston, 2006).

*Alliaria petiolata* (M. Bieb) Cavara and Grande (Brassicaceae), colloquially known as garlic mustard, is an herbaceous biennial plant native to Eurasia that has gained notoriety as invasive species for producing allelopathic biochemicals that disrupt the soil microbiome. Specifically, *A. petiolata* suppresses arbuscular mycorrhizal fungi (AMF) associated with native plants under controlled laboratory experiments (reviewed in Cipollini and Cipollini, 2016). AMF colonize the roots of most vascular plant species (Wang & Qiu, 2008), and provide nutrients to their symbiotic partners in exchange for essential carbohydrates (Smith and Read, 1996). Furthermore, the symbiosis can reduce rates of pathogen infection (Azcón-Aguilar, Jaizme-Vega, & Calvet, 2002; Borowicz, 2001; Smith & Read, 1996; Wehner, Antunes, Powell, Mazukatow & Rillig, 2010). As such, AMF can have a strong influence on above-ground plant community composition and productivity (Koyama, Maherali & Antunes, 2019; Klironomos, McCune, Hart & Neville, 2001; van der Heijden et al., 1998).

The Novel Weapons Hypothesis (NWH) states that some plants can invade by producing toxic biochemicals for which native competitors have not evolved countermeasures (Callaway & Ridenour, 2004). This hypothesis is supported by the observation that *A. petiolata* leaf and root extracts can inhibit AMF spore germination in laboratory assays (Koch et al., 2011; Roberts & Anderson, 2001; Stinson et al., 2006). Moreover, conditioning sterile soils with *A. petiolata* can reduce AMF root colonization across multiple hosts (Koch et al., 2011; Stinson et al., 2006). However, in contrast to controlled experiments, results from field studies provide mixed support for *A. petiolata* suppressing AMF in its introduced range. For example, *Arisaema triphyllum*, *Maianthemum racemosum* and *Trillium grandiflorum* growing in presence of *A. petiolata* had similar species richness of AMF and abundance of colonization in roots relative to their conspecifics growing outside of the invasion zone, and AMF composition was unaffected in *A. triphyllum* (Burke, 2008). In contrast, co-occurrence with *A. petiolata* reduced the extend of AMF colonization in *Acer saccharum* saplings even though AMF richness remained unaltered (Barto et al., 2011). Such inconsistency in the literature could be explained by plant-specific AMF-dependence (Stinson et al., 2006) or differences in experimental context. It is therefore unclear how soil AMF community structure is affected by natural *A. petiolata* populations.

Because of *A. petiolata*’s putative suppression of AMF, research on the effect of *A. petiolata* on soil microbial functional groups other than AMF has also generated interest. Lankau (2011a) measured overall fungal diversity in response to conditioning by *A. petiolata* from populations with varying concentration of glucosinolates and found that the highest richness of fungi occurred at intermediate concentrations. This response suggests that fungi have a complex relationship with *A. petiolata* and individual fungal taxa could be either stimulated or inhibited by *A. petiolata* depending on their functional role. Recently, Anthony, Frey, and Stinson (2017) compared the composition of different fungal functional groups between *A. petiolata* invaded and uninvaded soils using ITS metabarcoding. They found that the presence of *A. petiolata* reduced ectomycorrhizal fungi (EMF), but was associated with an increase in saprotrophic decomposers and fungal pathogens. These findings provide some support for the Enemy of my Enemy Hypothesis (EEH), which suggests that invasive species can outperform native species by accumulating pathogens that are more harmful to competitors (Colautti, Ricciardi, Grigorovich & MacIsaac, 2004; Flory & Clay, 2013). However, the effects of fungal community changes on plant health were not examined, and the forward primer fITS7 of the 5.8S rRNA gene is a poor match for AMF and other Glomeromycotina (Taylor et al., 2016). Thus, the relative effects of AMF versus non-mycorrhizal groups is a key aspect that remains unclear.

Contrary to the interest generated by fungi, the impact of *A. petiolata* on bacterial communities has received little attention and studies have generally used Terminal restriction fragment length polymorphism (TRFLP) profiles. Although TRFLP has the potential to compare broad trends in community composition, it may not provide enough taxonomic resolution to identify the impacts of *A. petiolata* invasion on soil microbiota and their functional roles. Based on TRFLP profiles, Lankau (2011b) found that geographically separated populations of *A. petiolata* had significantly distinct soil fungal and bacterial richness and community composition. However, there were no consistent differences in fungal and bacterial community composition or richness between invaded and adjacent (at least 20 m away from actively growing *A*. *petiolata* populations) uninvaded soils. Similarly, Burke and Chan (2010) found that *A. petiolata* had little to no effect on the bacterial species richness present in the topsoil when compared to soil found under the native herb *Allium tricoccum* and *Gallium triflorum*. However, they noted that bacterial community composition was different in soil collected under *A. petiolata* compared to soil collected under native herbs, but only during the month of August and at no other time during the growing season. Thus, despite some evidence that *A. petiolata* invasion may not affect bacterial richness, its effect on bacterial community composition and function remain unclear.

Here we test whether *A. petiolata* invasion is linked to feedbacks causing changes in structure and function of soil microbial communities as predicted by the EEH and NWH. We examine the extent to which such changes hinge on the suppression of mutualists (NWH) and the recruitment of pathogens (EEH). In addition, we assess AMF colonization and pathogen lesions on the roots of plant species that co-occur with *A. petiolata* in the field, combined with metabarcoding of microbial DNA sequences in soil samples. We then assigned functional groups to microbial DNA sequences to link shifts in microbial community to changes in plant health and ecosystem processes.

## Materials and Methods

### Site description

We conducted the study at the Queen’s University Biological Station (QUBS) located in the Frontenac Axis near Elgin, Ontario, Canada (44.5675° N, 76.3245° W). This is the first report of a study conducted at this site. The climate in the region is temperate, with the highest levels of precipitation occurring in October and November. We located ten naturally occurring populations of *Alliaria petiolata* containing a minimum of one hundred individuals of both first-year rosettes and second-year reproductive individuals. All ten populations were within 5 km of each other, but at least 150 m apart (see Supplementary material for a list of the site coordinates and maps). The forest canopies of all populations were dominated by *Acer saccharum*, *Fraxinus americana*, *Pinus stobus* and *Ostrya virginiana*, resulting in a moderately well shaded understory. Eight of the sites coincided with walking trails and old abandoned fields which increased the amount of disturbance. During the summer months, the understories were primarily composed of *Acer saccharum* saplings, *Anemone americana*, *Solidago canadensis*, and *Galium aparine*. We also found the presence of *Trillium grandiflorum* and *Claytonia virginica* in the early spring and *Ariseama triphyllum*, *Maianthemum racemosum*, and *Erythronium americanum* in the summer. The *A. petiolata* populations in this study were all found in podzol or brunisol soils, which are characterized by good drainage and mesic to sub-mesic moisture regimes. All populations had their soil texture classified as either a loam or a sandy loam. Most populations were found on the crest of or along mild slopped hills. In all the sites, we found signs of *Lumbricus terrestris* invasion, which was expected since the removal of *A. petiolata* has been shown to reduce *L. terrestris* densities (Stinson et al. 2018).

### Site establishment and sample collection

We established transects through each population, running from east to west, except where there was a steep incline. In those cases, we changed the orientation of the transect to minimize differences in incline between the middle and the end of the transect. We ended transects 7 m away from the edge of the population, which was determined as being where the last visible *A. petiolata* plant was present along or 7 m around the transect.

We identified six species that consistently co-occurred with *A. petiolata* and are known to interact with AMF: *Geranium robertianum*, *H. americana*, *S. canadensis*, *Circaea lutetiana*, *G. aparine*, and *M. racemosum*. All species are listed as native in Vascan (Brouillet et al. 2010). The USDA database also lists *H. americana*, *S. canadensis, M. racemosum, G. robertianum* and *C. lutetiana* as native to Ontario (USDA, NRCS, 2018). However, *G. aparine* is listed as uncertain in Ontario but native in New York and *G. robertianum* is listed as native in Ontario but uncertain in New York on the USDA database (USDA, NRCS, 2018). Therefore, although the status of *G. robertianum* and *G. aparine* are uncertain, they seem to be Holarctic native or pre-colonial introductions.

In the last week of June 2016, we sampled soil and the roots of the plants identified above in all populations. To characterize the edaphic properties, we used an auger (5 by 5 cm) to collect 45 mL of topsoil where the highest density of root and fungal hyphae can be found. We collected soil in triplicate, 1m apart, around both the middle (henceforth ‘invaded’) and the end (henceforth ‘uninvaded’) of the transect through the actively growing *A. petiolata* population for a total of six samples per population. We sterilized the tools with 70% ethanol in-between sample collections. Following collection, the triplicate soil samples were pooled and stored at −20°C. We collected the roots of six specimens per plant species when we could find them in a population: three inside, and three outside the *A. petiolata* populations. The plants collected outside were found at a minimum of 7 m from the closest *A. petiolata* individual at the edge of the population. We chose to use 7 m as the minimum distance from the edge of a population to limit potential effects of *A. petiolata* on the uninvaded soil, while limiting differences in abiotic soil characteristics (Rodgers, Wolfe, Werden & Finzi, 2008; Wolfe, Rodgers, Stinson & Pringle, 2008). To collect the root, we gently dug approximately 5 cm around the bottom of the plant and lifted the individual from under its root system. We took the whole root system from the ground with the surrounding soil. Once removed from the ground, we shook off the soil from the roots and took the samples to be analyzed. In the laboratory, individual root samples were washed in distilled water and preserved in 50 mL falcon tubes immersed in 70% ethanol.

### Soil physical properties

We analyzed soil samples for soil characteristics that influence AMF species diversity (Lauber, Strickland, Bradford & Fierer, 2008). Total carbon and nitrogen were measured using a subsample of 60 mg (dry mass) of oven-dried soil in the Flash 2000 Combustion NC Soil Analyzer following the analyzer’s protocol (Thermo Fisher, Waltham, MA, USA) with 10 mg of aspartic acid as control. We vortexed a mixture of 20 mL of water with 10 g of soil every 15 minutes for an hour and measured the pH with a potentiometric reader (Thermo Fisher, Waltham, MA, USA). To measure soil aggregate stability, we processed 4 g of 2 mm airdried aggregates in the Eijkelkamp sieving apparatus using the apparatus’ protocol (Eijkelkamp, Giesbeek, The Netherlands). We measured the soil texture using the Hydrometer method (Thermo Fisher, Waltham, MA, USA) with a 10% sodium hexametaphosphate dispersing agent. Further details on our protocols are provided as Supplementary Materials.

### Arbuscular mycorrhizal fungi and lesion root colonization

We stained native plant roots using a modified version of the protocol developed by Vierheilig, Coughlan, Wyss, and Piché (1998) which uses Sheaffer black ink to visualize AMF colonization. Briefly, we cut the roots of each plant into 2 cm fragments and placed them in a labeled tissue cassette (Starplex Scientific Corp, Cleveland, TN, USA). We placed the cassette in KOH solution and left it in a water bath at 90 °C for two hours before cleaning it with deionized water and re-acidifying it in 10 % acetic acid for 10 minutes. We then moved the cassette to the ink vinegar solution in a beaker in the 90 °C water bath for a total for 7 minutes. Finally, we rinsed the cassette in deionized water and immersed it in lactoglycerol for a minimum of 20 minutes to preserve the roots before preparing permanent slides.

We assessed root AMF colonization following methodology by McGonigle, Miller, Evans, Fairchild, and J. Swan (1990). Briefly, we scored arbuscules, vesicles, or AMF hyphae under a microscope at 100 roots intersections with the microscope’s field of view (200x) for each individual sample. We scored the AMF traits in a hierarchical manner, if AMF hyphae were present, we scored hyphae, but if vesicles were present, we only scored vesicles, and if arbuscules were present, we only scored arbuscules. If AMF were absent this was recorded too. This method resulted in a binary score for signs of AMF at each cross-section (henceforth ‘AMF colonization’). To score lesions, we used a modified version of this protocol to identify potential pathogenic effects by searching for signs of decay, and non-AMF hyphae (henceforth lesions). Reference images of lesion or mycorrhiza refer to Supplementary Materials Figure S2 and S3.

### DNA extraction, library construction, and sequencing

We extracted DNA from 10 g of pooled soil using a PowerMax Soil DNA Isolation Kit (MoBio Laboratories Inc., Carlsbad, California). We assessed the quality and concentration of purified DNA using a NanoDrop Spectrophotometer (NanoDrop Technologies, Wilmington, DE, USA). We used a two-step PCR amplification, with the first PCR reaction in triplicate following the method developed by Fadrosh, Gajer, Sengamalay, Brotman and Ravel (2014). The first PCR reaction aimed to isolate and amplify the template DNA in the genomic DNA for each sample. After the first reaction, we cleaned and pooled the product of the three replicates for the second PCR reaction, which used barcoded primers (Wu et al. 2015). The separation between the first and second PCR reaction was used to reduce any potential bias introduced by the large, barcoded primers.

We performed targeted amplification of the bacterial 16S ribosomal-RNA region and the fungal internal transcribed spacer regions (ITS) to characterize the composition of soil microbial communities. To amplify bacteria and archaea in the soil samples we analyzed bacterial communities by targeting variable region 4 of the 16S ribosomal gene using the improved 16S rRNA Gene primer recommended by the Earth Microbiome Project: 515f Modified [5′-GTGYCAGCMGCCGCGGTAA-3′] and 806r Modified [5′-GGACTACNVGGGTWTCTAAT-3′] (Walters et al., 2015). To amplify sequences from fungal organisms, we chose to use ITS region because it is better at discriminating between microbial species and assigning functional groups (Schoch et al., 2012). Specifically, we used Taylor et al.’s (2016) 5.8S-Fun [5′-AACTTTYRRCAAYGGATCWCT-3′] and ITS4-Fun [5′-AGCCTCCGCTTATTGATATGCTTAART-3′] primer pair (hereafter simply ITS) because, unlike other primers targeting the ITS region, this primer pair minimizes the amplification of non-target eukaryotes to maximize the coverage for the target fungi. Additionally, these primers have been successfully used in other studies to identify AMF species (Gao et al., 2019; Krüger, Krüger, Stockinger & Schüßler, 2012). We processed each sample with two technical replicates per primer to ensure the validity of the molecular methods.

The PCR method we used contained two reactions. The first reaction amplified the 16S or ITS region, and the second reaction attached the larger barcode, spacer, and Illumina sequencing primer region. The first PCR step was a 50 μL reaction which contained the following reagents: 0.25 mM forward and 0.25 mM reverse target-only primers (Eurofins Scientific, Brussels, Belgium), 1.25 mM MgCl (Invitrogen, Carlsbad, CA, USA), 0.04 mM BSA (Ambion, Foster City, CA, USA), 0.1 mM dNTPs (Invitrogen, Carlsbad, CA, USA), 0.02 U/μL of Platinum™ Taq DNA Polymerase High Fidelity (Invitrogen, Carlsbad, CA, USA), 2.5 μL of 10x PCR buffer, which was composed of 100mM Tris HCl at pH 8.4 and 25 mM KCl (Invitrogen, Carlsbad, CA, USA), and finally 0.6 ng/μL of the genomic DNA. This first reaction amplified the 16S or the ITS targeted segments from the template DNA for 12 cycles with a denaturing step of 96°C for 30 seconds, an annealing step of 58°C for 30 seconds, and an elongation step of 72°C for 1 minute and 30 seconds.

Following the first PCR, we purified the amplified DNA using 1.8x AMPure beads (Beckman Coulter, Beverly, MA, USA). The purified PCR product was used as template for a second PCR step, which was a 25 μL reaction with 0.4 mM forward and 0.4 mM reverse primers, 10 mM Tris HCl at pH 8.4, 25 mM KCl, 2.5 mM MgCl, 0.08 nM BSA, 0.2 mM dNTPs, and 0.04 U/μL of Platinum™ Taq DNA Polymerase High Fidelity. We used 15 μL of the purified PCR product of step one as the template DNA of step 2. We amplified the 16S or ITS segment for 28 cycles with the same temperature and time specifications as the first PCR reaction. We cleaned the final product with the 1.8x AMPure bead mixture. We measured the molarity of the samples using PicoGreen reagents on the Qubit Fluorometer (Life Technologies, Grand Island, NY, USA), normalized, and pooled all samples, including both ITS and 16S, to a final concentration of 4 pM in EDTA. The library, including ITS and 16S amplicons, was sequenced on a single lane of the MiSeq platform at Queen’s University (Kingston, Ontario, Canada) using the MiSeq Reagent Kit v3, with 600 cycles (Illumina, San Diego, CA, USA). The raw sequences were deposited in the NCBI Short-Read Archive (BioProject PRJNA551217).

### Bioinformatic pipeline

Raw sequences were first quality filtered and sorted using the process_radtags command from the Stacks software package (Catchen, Hohenlohe, Bassham, Amores & Cresko, 2013) to demultiplex the barcoded sequences, allowing for two mismatches in the barcode sequence and disabling the restriction site associated DNA marker check. They were then sorted by primer to differentiate 16S and ITS amplicons using the seal.sh command from BBmap (Bushnell 2016) with 5 base-pair-long k-mers. The demultiplexed sequences were imported into QIIME2 (Bolyen et al., 2019), and using the DADA2 plug-in (Callahan et al., 2016) we merged paired-end reads, filtered chimeric reads, and produced a table of amplicon sequence variants (ASV). We assigned taxonomy to each ASV using BLAST+ (Camacho et al., 2009) with the Silva database version 1.32 (Quast et al., 2013) for 16S samples and the UNITE database version 7.2 (Kõljalg et al., 2013) for ITS samples. A sequence was assigned taxonomy if ≥99% of the sequence aligned to a reference and matched the ASV with ≥97% identity (Schloss & Westcott, 2011). We allowed for 1 BLAST hit, and 51% of the ASV had to match the top hit with 100% precision to accept it as the consensus taxonomy assignment. After quality control and ASV assignment we retained 9 325 402 sequences, which were assigned to 6 907 ASVs across both bacteria and fungi. We submitted the sequences to the NCBI Sequence Read Archive (SRA) under the BioProject PRJNA551217.

The Funguild program annotates the species composition table with a functional guild (e.g. AMF, EMF, dung saprotroph, wood saprotroph) for each assigned fungal taxa (Nguyen et al., 2016). Following taxonomic assignment, we used the program to group fungal species from our samples into five functional groups: AMF, EMF, saprobes (including wood saprotroph, soil saprotroph, and undefined saprotroph), fungal animal pathogens, and fungal plant pathogens. Funguild notes that functional roles are often context dependent, thus their guild assignment for fungi that have multiple groups have a lower confidence ranking. Pathogenicity is particularly context dependent and, as such, we kept and considered all Funguild assignments to the pathogen group despite their lower confidence ranking. Consequently, we consider all references to pathogens as putative pathogens. There is no tool similar to Funguild available for soil bacteria; thus, we manually researched indicator species for the invaded and uninvaded soils (**see Indicator species analysis section**) using citations from Google Scholar and Web of Science based on their phylogenic classification. In the statistical program R version 3.3.2 (R core team, 2018), we preprocessed our samples by removing ASV that occurred only once or were unassigned at the phylum level (842 fungi ASVs and 315 bacteria ASVs) using the package *phyloseq* (McMurdie & Holmes, 2013). The bioinformatic pipeline was deposited in DRYAD (doi: 10.5061/dryad.hqbzkh1bk).

### Statistical analysis

All statistical analyses were done in the statistical program R version 3.3.2 (R core team, 2018). For all statistical tests in this analysis, we identified the model of best fit using likelihood ratio tests of hierarchical models with the *anova* function in R and removing parameters that were not significant predictors (P < 0.05). We deposited the reproducible data analysis in DRYAD as Markdown-formatted text converted to HTML (doi: 10.5061/dryad.hqbzkh1bk).

Differences in edaphic factors associated with *A. petiolata* presence could be a cause or consequence of invasion. To determine which soil factors should be included as predictors in our microbial composition models, we tested which edaphic factors were significantly associated with presence of *A. petiolata* using a binomial generalized linear model with invasion status as the response variable. Aggregate stability and pH were included as response variables, along with a and a principal component axis representing carbon and nitrogen concentration. We grouped carbon and nitrogen concentration data into a single PC axis due to high collinearity (*r* = 0.95). Only pH was significantly associated with *A. petiolata* presence and retained for the analysis detailed below.

We used multivariate ordination to characterize microbial communities and functional groups, and then used generalized linear models and permutation to test the effects of *A. petiolata* and edaphic parameters on microbial composition. We used the ***adonis2*** function in the *vegan* package (Oksanen et al., 2018) with 999 permutations to test the response of the relative abundance of bacteria and fungi to invasion status, population, technical replicate, and pH. The other edaphic variables were not significantly associated with *A. petiolata* presence, as such, we will not be including them in further regression models to avoid overfitting them. To visualize differences in species composition, we plotted non-metric multidimensional scaling (NMDS) coordinates for two dimensions based on Bray-Curtis dissimilarity matrices using ***metaMDS*** from *vegan*. We fit a generalized linear mixed model, as implemented by the ***glmmadmb*** function from the *glmmADMB* package (Fournier et al., 2016) to test for the effect of invasion and pH on bacterial and fungal species richness using species richness as the response variable with *A. petiolata* invasion status as the explanatory variable, and technical replicate nested within population as a random factor. When beta-diversity significantly differed between invaded and uninvaded soils we used the ***beta.pair*** function from the *betapart* package (Baselga & Orme, 2018) with 999 permutations to partitioned beta diversity into spatial turnover, indicating species replacement with no reduction in diversity, and nestedness, indicating species loss and reduced diversity. We tested the significance of the correlation of invasion with turnover or nestedness of species composition using Multiple Response Permutation Procedure (MRPP) with *vegan*’s ***mrpp*** with 999 permutations. We analyzed the effect of *A. petiolata* on the composition of the five functional groups of fungi following the same statistical analysis outlined for overall fungal and bacterial composition.

To determine how changes in soil microbial communities impacted the root health of co-occurring plant species, we tested the effect of *A. petiolata* presence on the proportion roots colonized by AMF and the proportion roots affected by lesions. Because the distribution of the proportions was normal and included both zeros and ones, we used a linear mixed effects model as implemented by ***lmer*** function from the *lme4* package in R (Bates et al., 2015). Specifically, we tested the either the proportion of arbuscular mycorrhiza or the proportion of lesions on roots of each individual plant as a response variable. Predictor variables for each model included host species, invasion, and the interaction of host species and invasion, with population as a random effect.

### Indicator species analysis

We tested whether soil invaded by *A*. *petiolata* or soil near *A*. *petiolata* invasion had nonrandom associations with bacterial ASV using the indicator species analysis (Dufrêne & Legendre, 1997). The indicator species index varies from 0 to 1 and is a measurement of the strength of association of the taxa to its assigned group based on fidelity and relative abundances of the ASV. To implement this analysis, we used the ***multipatt*** command from *indicspecies* package (De Cáceres & Legendre, 2009), which tests for the significance of the indicator species index through a permutation test with 999 permutations. We researched the ecological function of each taxa that differed between invaded and uninvaded areas separately using the taxonomic assignment of the ASV.

In addition to identifying indicator taxa, we wanted to determine if particular functional groups were suppressed or stimulated by *A. petiolata* invasion, under the null model that indicator taxa are a random sample of available functional groups. We analyzed stimulated and suppressed indicator species separately and randomly sampled the total number of ASVs from each group, with 1000 iteration each. This produced a null distribution of the frequency of each functional group. This approach is analogous to a gene set enrichment analysis of gene expression data (Subramanian et al., 2005). The indicator species analysis is available as part of the reproducible data analysis in DRYAD (doi: 10.5061/dryad.hqbzkh1bk).

## Results

### Edaphic properties

Most edaphic properties in the core of *Alliaria petiolata* populations did not differ significantly from those outside but near the populations. In fact, texture (n = 20, Φ = 1.77, Deviance df: 2 = 0.87, p-value > 0.05), aggregate stability (n = 20, Φ = 1.77, Deviance df: 1 = 1.17, p-value > 0.05), and a principal component accounting for carbon and nitrogen composition (n = 20, Φ = 1.77, Deviance df: 1= 0.02, p-value > 0.05) were the same for soil samples collected inside and outside the *A. petiolata* invasion zone. However, pH significantly (n = 20, Φ = 1.77, Deviance df: 1= 4.05, p-value < 0.05) decreased outside of the invaded area (Estimate=−0.82, CI= −1.98, 0.13) (**Fig. 1**).

**Figure 1.**
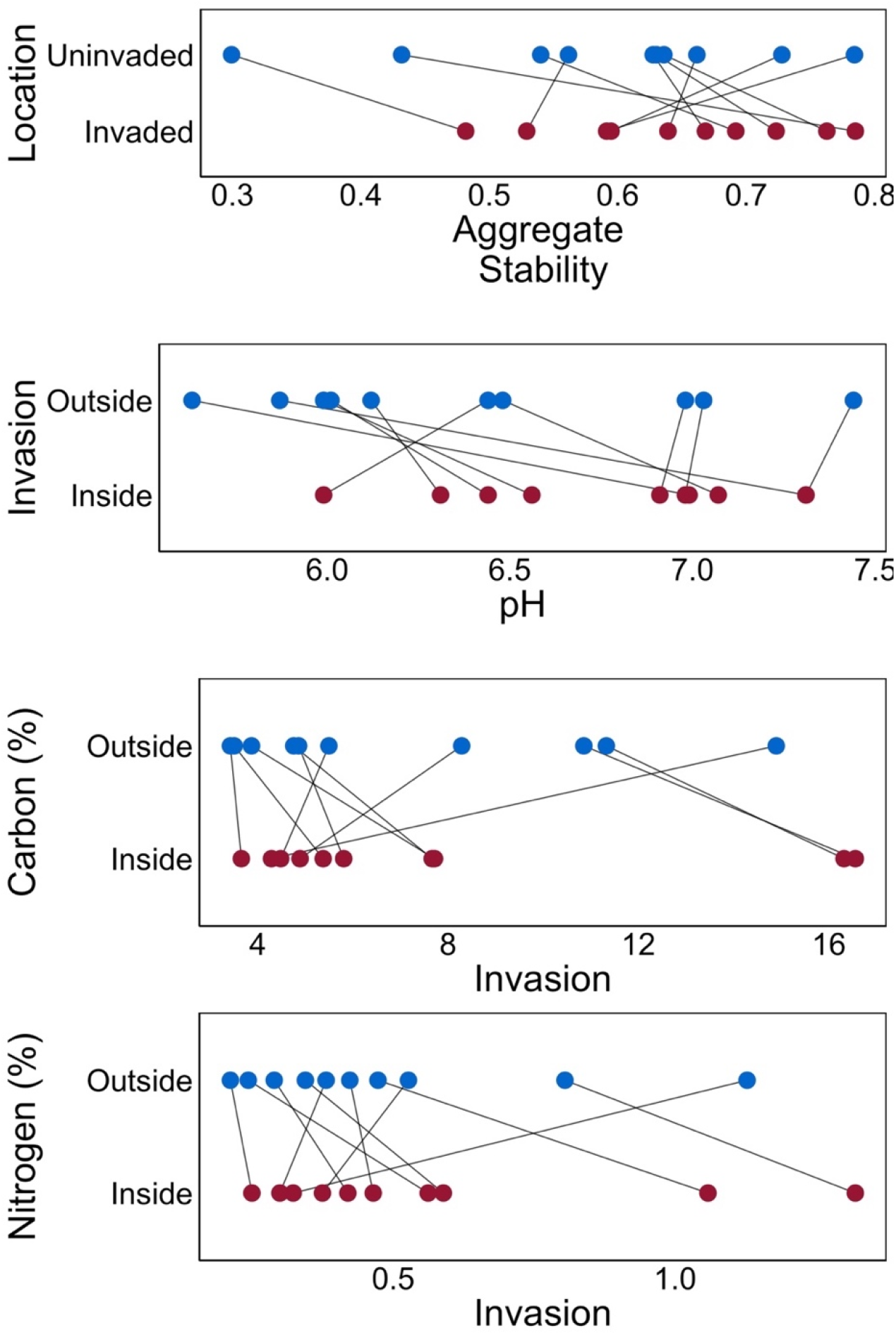
Soil physical properties of soil properties inside and outside the invasion zone. Points showing the value of each soil physical property either inside the A. petiolata invaded area (inside) or outside the invaded area (outside). The values inside and outside the *A. petiolata* invasion for each population are linked by a gray line. The plots show aggregate stability, pH, percent carbon and percent nitrogen content of the soil. Only pH differed significantly between invaded and uninvaded soil (P < 0.05).

### Diversity and community composition

The most abundant fungal phyla were Ascomycota (2804 ASV), Basidiomycota (1901 ASV), and Mortierellomycota (375 ASV). The most abundant fungal species present in soil were *Mortierella alpine* (40 samples), and *Mortierella minutissima* (40 samples), both saprobes. Additional information about phylum, family, and functional ASV composition is available as Supplemental information. Fungal community structure was different between invaded and uninvaded soils (n = 40, Deviance _d.f.:1_ = 0.57, R^2^= 0.030, p-value < 0.001), among populations (n = 40, Deviance_d.f.:9_ = 4.84, R^2^= 0.252, p-value < 0.001), and between technical replicates (n = 40, Deviance_d.f.:1_ = 0.52, R^2^= 0.027, p-value < 0.05). The differences in pH between invaded and uninvaded soils correlated significantly with pH on fungal community composition (n = 40, Deviance_d.f.:1_ = 0.54, R^2^= 0.028, p-value < 0.001). When beta diversity was divided into spatial turnover and nestedness, only spatial turnover responded to invasion (n _in,out_ = 20, 20, delta _in,out_ = 0.9932, 0.9938, p-value < 0.05), not nestedness (n _in,out_ = 20, 20, delta _in,out_ = 0.001, 0.001424, p-value > 0.05). Fungal species richness significantly (n = 40, Deviance_d.f.:1_ = 10.35, p-value < 0.001) increased inside *A. petiolata* invaded soil (CI: 23.42 to 85.33, Effect size: 54.37) (**Fig. 2**).

**Figure 2.**
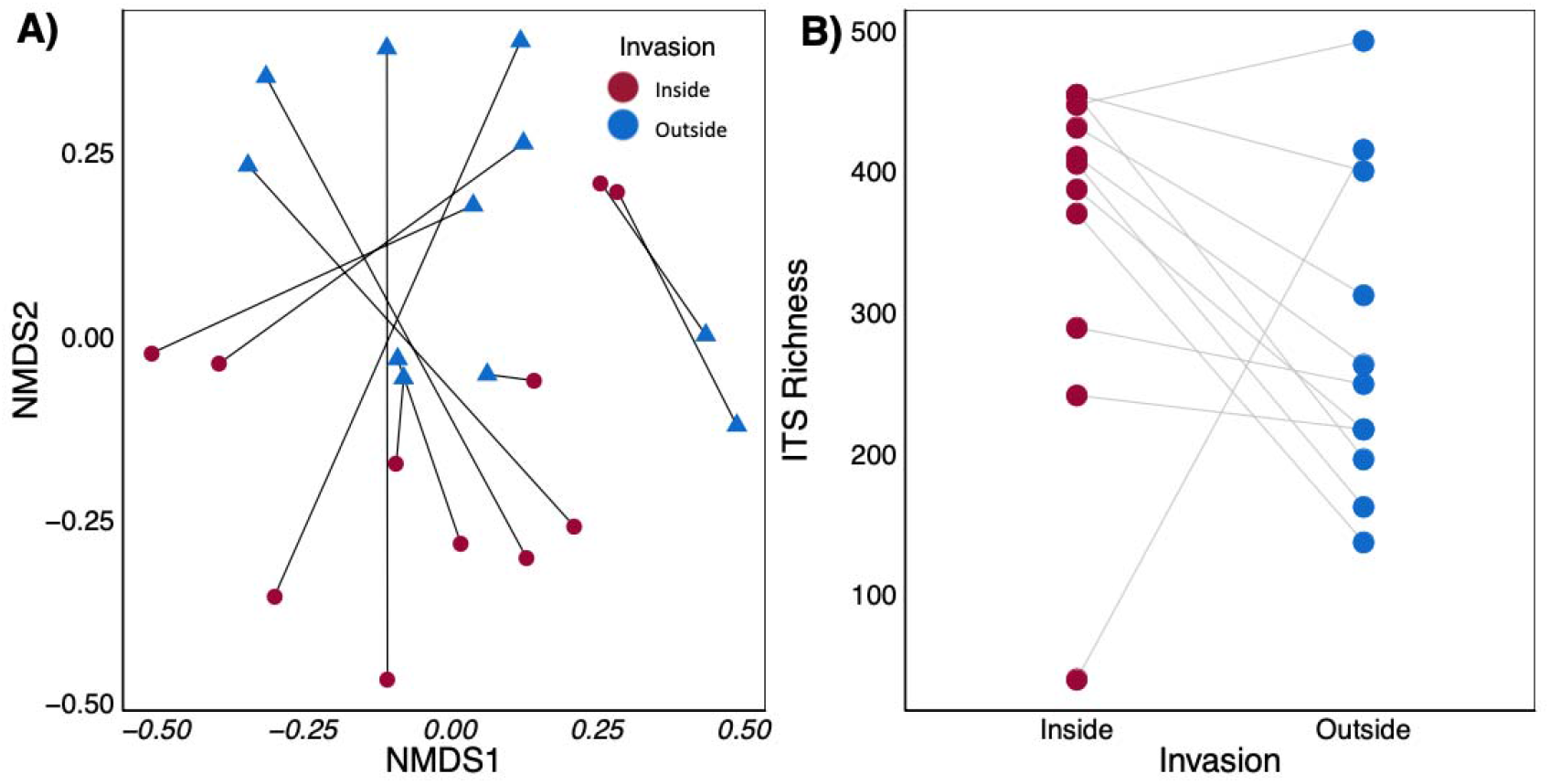
Invasion and fungal population community and richness. **A**) NMDS ordination of fungal community composition in soils using a Bray-Curtis distance matrix. Each point represents the mean species composition of two technical replicates of each location (inside vs. outside) in each population. The inside and outside samples from each of the *A. petiolata* populations are linked by a black line. NMDS axes stress = 0.23 after 64 iterations. Fungal community composition was significantly affected by invasion (P < 0.001) and population (P < 0.001). **B**) Plot with points representing fungal richness (as measured by number of ASV per sample) inside and outside of each *A. petiolata* population. Inside and outside points of each population are linked by a grey line. Fungal richness was significantly affected by invasion (P < 0.001)

The most abundant phyla identified in the 16S analysis were *Proteobacteria* (1388 ASV), followed by *Actinobacteria* (955 ASV) and *Acidobacteria* (779 ASV). Bacterial community structure differed with invasion (n = 40, Deviance_d.f.:1_ = 0.53, R^2^= 0.028, p-value < 0.01) and *A. petiolata* population (n = 40, Deviance_d.f.:9_ = 4.65, R^2^= 0.25, p-value < 0.001) (**Fig. 3**). Bacterial community composition was unaffected by pH (n = 40, Deviance _d.f.:1_ = 0.47, R^2^= 0.025, p-value 0.05) but differed significantly among technical replicates (n = 40, Deviance _d.f.:1_ = 0.52, R^2^= 0.028, p-value < 0.05). The change in bacterial species composition that occurred in the invaded sites compared to the uninvaded sites was related to species turnover (n _in,out_ = 20, 20, delta _in,out_ = 0.977, 0.969, p-value < 0.01) rather than a loss of species (n _in,out_ = 20, 20, delta _in,out_ = 0.006, 0.007, p-value > 0.05). Bacterial species richness was similar (CI: −36.194 to 32.957, Effect size: −1.619) inside as compared to outside *A. petiolata* invasion (n = 40, Deviance_d.f.:1_ = 0.008, p-value > 0.05).

**Figure 3.**
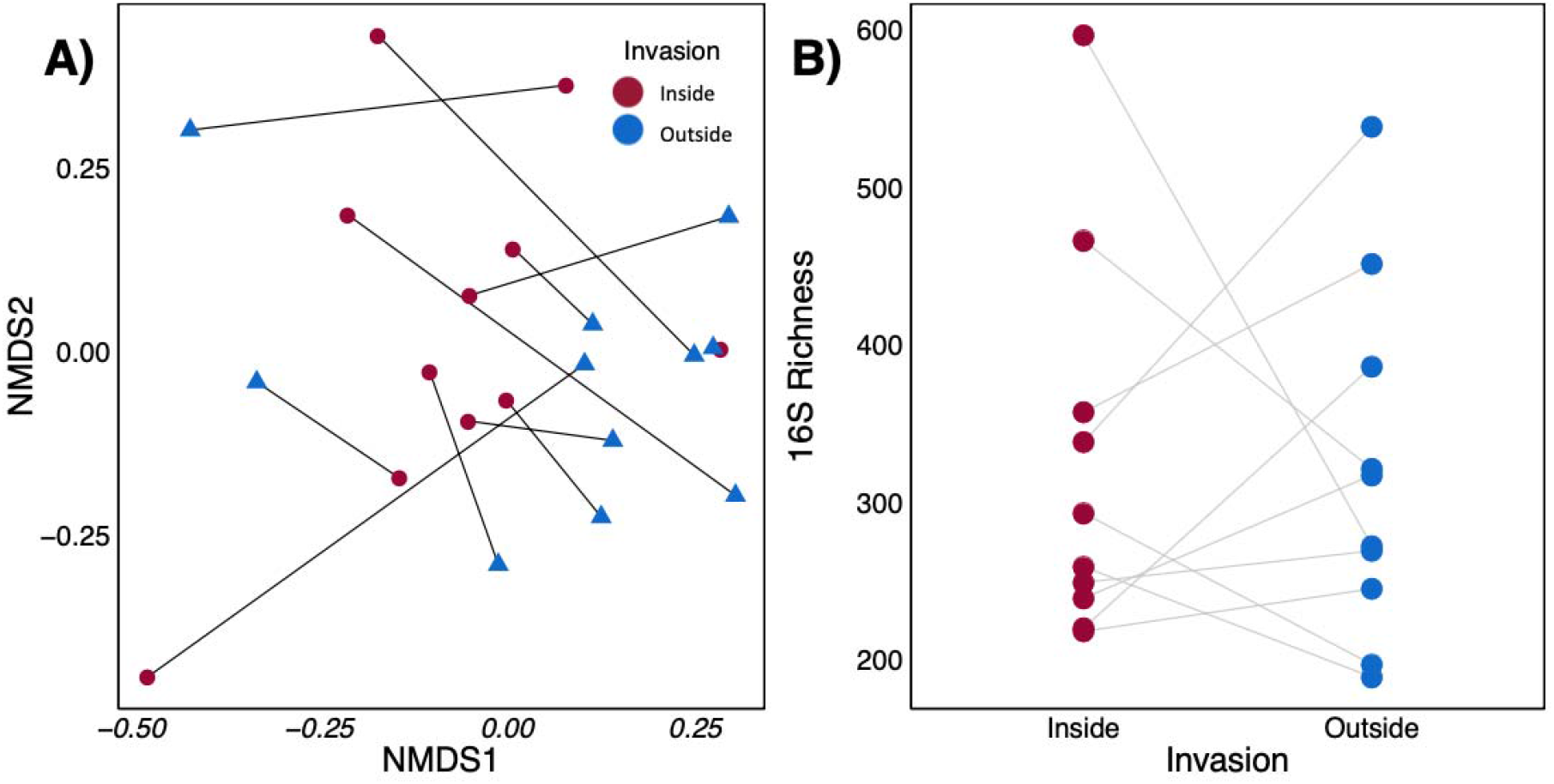
Invasion and bacterial population community and richness. **A**) NMDS ordination of Bacterial and Archaeal community composition in soils using a Bray-Curtis distance matrix. Each point represents the mean species composition of two technical replicates in each population (inside vs. outside). The inside and outside samples from each *A. petiolata* population are linked by a black line. NMDS axes stress = 0.19 after 20 iterations. Bacterial community composition was significantly affected by invasion (P < 0.01) and population (P < 0.001). **B**) Plot with points representing Bacterial and Archaeal richness (as measured by number of ASV per sample) inside and outside each *A. petiolata* population. Inside and outside points of each population are linked by a grey line. Bacterial richness was not significantly affected by invasion (P > 0.05)

### Changes in fungal functional groups

Arbuscular mycorrhizal fungi (179 ASVs) were heavily dominated in abundance by Glomeraceae (138 ASVs) but also included members of the families Claroideoglomeraceae (12 ASVs), Diversisporaceae (4 ASVs), Acaulosporaceae (2 ASVs) and Paraglomeraceae (1 ASVs). AMF community composition was similar in all populations (n = 33, Deviance _d.f.:9_ = 4.498, R^2^= 0.281, p-value > 0.05) and in invaded and uninvaded areas (n = 33, Deviance_d.f.:1_ = 0.502, R^2^= 0.031, p-value > 0.05). There were no significant differences among technical replicates for AMF (n = 33, Deviance_d.f.:39_ = 0.503, R^2^= 0.031, p-value > 0.05) or pH (n = 33, Deviance_d.f.:1_ = 0.493, R^2^= 0.031, p-value > 0.05). Additionally, invasion did not significantly relate to the richness of AMF (n = 40, Deviance_d.f.:1_ = 2.238, p-value > 0.05; CI: −0.734 to 0.088, Effect size: −0.323).

Overall, EMF (672 ASVs) were dominated in abundance by Thelephoraceae (23 ASVs), Inocybaceae (21 ASVs), and Hymenogastraceae (3 ASVs). The EMF community composition varied significantly among *A. petiolata* populations (n = 40, Deviance_d.f.:9_ = 4.84, R^2^= 0.250, p-value < 0.001) and differed between invaded and uninvaded soils (n = 40, Deviance_d.f.:1_ = 0.54, R^2^= 0.028, p-value < 0.01). There was no significant effect of technical replicate (n = 40, Deviance_d.f.:39_ = 0.46, R^2^= 0.024, p-value > 0.05) but pH significantly related to fungal composition (n = 40, Deviance_d.f.:1_ = 0.55, R^2^= 0.028, p-value < 0.01) for EMF. Using beta-diversity partitioning, we could not determine conclusively if the change in EMF community composition was attributed to a turnover in species composition (n _in,out_ = 20, 20, delta _in,out_ = 0.98, 0.96, p-value > 0.05) or species nestedness (n _in,out_ = 20, 20, delta _in,out_ = 0.01, 0.02, p-value > 0.05). Nevertheless, invasion did not relate to the richness of EMF (n = 40, Deviance_d.f.:1_ = 0.211, p-value > 0.05; CI: −0.25, 0.16, Effect size: −0.048); thus it is more likely that EMF species composition changes occur because of the replacement rather than species loss.

Animal pathogen communities (149 ASVs), which were dominated in abundance by Chaetomiaceae (11 ASVs), Clavicipitaceae (10 ASVs), Cordycipitaceae (5 ASVs). Animal pathogens varied with *A. petiolata* invasion (n = 38, Deviance _d.f.:1_ = 0.687, R^2^= 0.038, p-value < 0.001) and with pH (n = 38, Deviance _d.f.:1_ = 0.63, R^2^= 0.035, p-value < 0.01), but not among populations (n = 38, Deviance _d.f.:9_ = 4.52, R^2^= 0.250, p-value > 0.05), or technical replicate (n = 38, Deviance_d.f.:37_ = 0.400, R^2^= 0.022, p-value > 0.05). We could not conclusively assign this change in community composition to species turnover or nestedness. However, animal pathogen richness significantly decreased (n = 40, Deviance_d.f.:1_ = 9.756, p-value < 0.01; CI: −5.22, −1.52, Effect size: −3.37) outside *A. petiolata* invasion.

Plant pathogens (566 ASVs) were dominated in abundance by Spizellomycetaceae (12 ASVs), Nectriaceae (14 ASVs), and Mortierellaceae (23 ASVs). Plant pathogen composition varied among populations (n = 40, Deviance _d.f.:9_ =4.72, R^2^= 0.245, p-value = 0.002) but not with invasion (n = 38, Deviance_d.f.:1_ = 0.532, R^2^= 0.028, p-value > 0.05), technical replicate variation (n = 38, Deviance_d.f.:37_ = 0.506, R^2^= 0.026, p-value = 0.193), or pH (n = 38, Deviance_d.f.:1_ = 0.514, R^2^= 0.027, p-value > 0.05),. However, plant pathogen richness was significantly different between *A. petiolata* invaded and *A. petiolata*-free areas (n = 40, Deviance _d.f.:1_ = 5.044, P < 0.05). That is, pathogen richness decreased (CI: −12.94, −1.11, Effect size: −7.02) outside of *A. petiolata* invaded soils.

The saprotroph community (2509 ASVs) was dominated in abundance by Mortierellaceae (171 ASVs) and Hygrophoraceae (39 ASVs), and Thelephoraceae (63 ASVs) and differed among *A. petiolata* invaded plots when compared to uninvaded plots (n = 40, Deviance _d.f.:1_ = 0.581, R^2^= 0.030, p-value < 0.001), as well as among populations (n = 40, Deviance _d.f.:9_ = 4.752, R^2^= 0.248, p-value < 0.001). Soil pH (n = 40, Deviance_d.f.:1_ = 0.52, R^2^= 0.027, p-value < 0.05) had an effect on saprotroph composition, but not technical replicate (n = 40, Deviance_d.f.:1_ = 0.473, R^2^= 0.025, p-value > 0.05). The community change of saprobes was a result of spatial turnover (n_in,out_ = 20, 20, delta _in,out_ = 0.97, 0.98, p-value < 0.01) rather than species loss (n_in,out_ = 20, 20, delta _in,out_ = 0.02, 0.02, p-value > 0.05). Additionally, saprobe richness significantly decreased (CI: −43.72 to −15.06, Effect Size: −29.39) outside of *A. petiolata* invasion (n = 40, Deviance _d.f.:3, 4_ = 13.152, p-value < 0.001) (**Fig. 4**). Detailed information on the taxonomic composition of each functional group is provided as Supplementary information.

**Figure 4.**
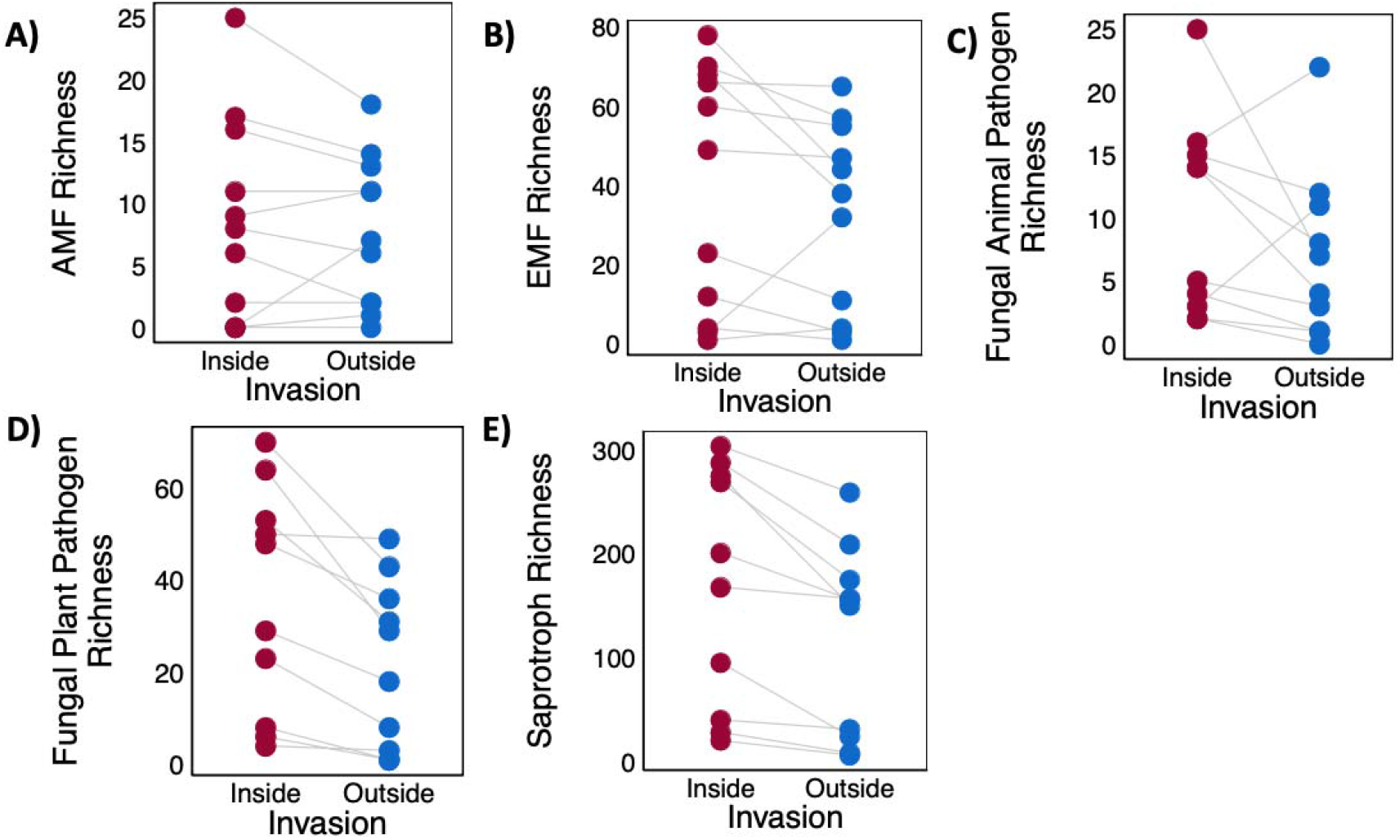
The richness for inside and outside the *A. petiolata* populations for fungal functional groups. Points representing average richness (as measured by number of ASV per sample) of two technical replicates collected inside and outside *A. petiolata* populations for each functional group: **A)** AMF, **B)** EMF, **C)** fungal animal pathogens, **D)** fungal plant pathogens, and **E)** saprotrophs. Inside and outside points for each population are linked by a grey line. Although community composition of EMF (P < 0.05), animal pathogens (P = 0.002), plant pathogens (P > 0.05), and saprotrophs (P < 0.001) differed with *A. petiolata* invasion, only animal pathogens (P 0.01), plant pathogen (P < 0.05), and saprotroph (P < 0.01) richness were significantly different inside and outside *A. petiolata* populations.

### Changes in bacterial functions

Overall, the most abundant families of bacteria overall were *Solirubrobacterales* 67-14 (104 ASVs), *Chthoniobacteraceae* (88 ASVs), and an unknown family of the phylum *Acidobacteria* in Subgroup 6 (280 ASVs). A total of 12 bacteria were indicator species for either invaded or uninvaded areas (**Table 1, Fig. 5**). Seven species were significantly associated with invaded soils and five with uninvaded soils. Because of the paucity of knowledge on bacterial function, not all bacteria that were identified as indicators could be assigned a function. Nonetheless, we were able to determine that of the five species associated with uninvaded soil three were associated with the nitrogen cycle. Specifically, we identified two nitrogen fixers of the family *Xanthobacteraceae* and one ammonia oxidizer of the family *Nitrosomonadaceae*. Based on enrichment analysis (see Supplemental information), amplicon sequence variants that functioned as nitrogen fixers were indicators of uninvaded soil samples more than expected by chance (p-value < 0.005). Of the seven species associated with invaded soil, two were part of the order *Gaiellales*. There was a trend toward *Gaiellales* (p-value > 0.05) being stimulated in soil invaded by *A. petiolata.*

**Figure 5.**
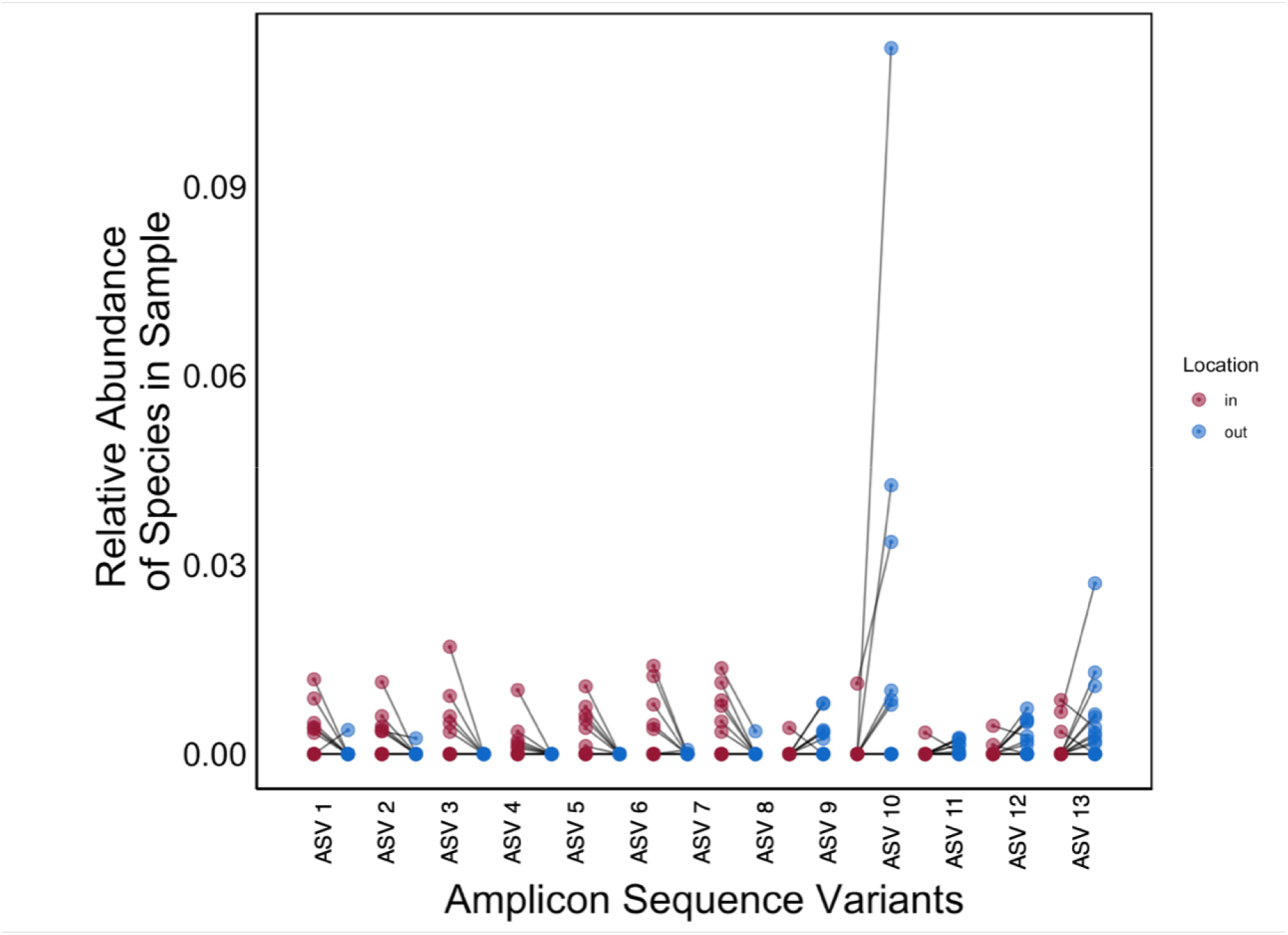
The relative abundance of amplicon sequence variants of bacteria and archaea that are indicators of samples from soils collected inside or outside *A. petiolata* populations, grouped by invaded (in) and non-invaded soil (out). The y-axis shows the relative abundance of each bacterial species that are specifically associated with samples from either inside or outside *A. petiolata* populations, grouped by *A. petiolata* invaded (inside, red) and *A. petiolata*-free (outside, blue) areas. Each point represents the relative abundance of the species in the sample taken from a population, either inside or outside. The values for inside and outside of a population are linked by grey lines. Amplicon sequence variant (ASV) numbers are the same as in column 1 of Table 1, which contains the phylogeny, the indicator value, the specificity, the fidelity, and functional information of each taxonomic group.

**Table 1.**
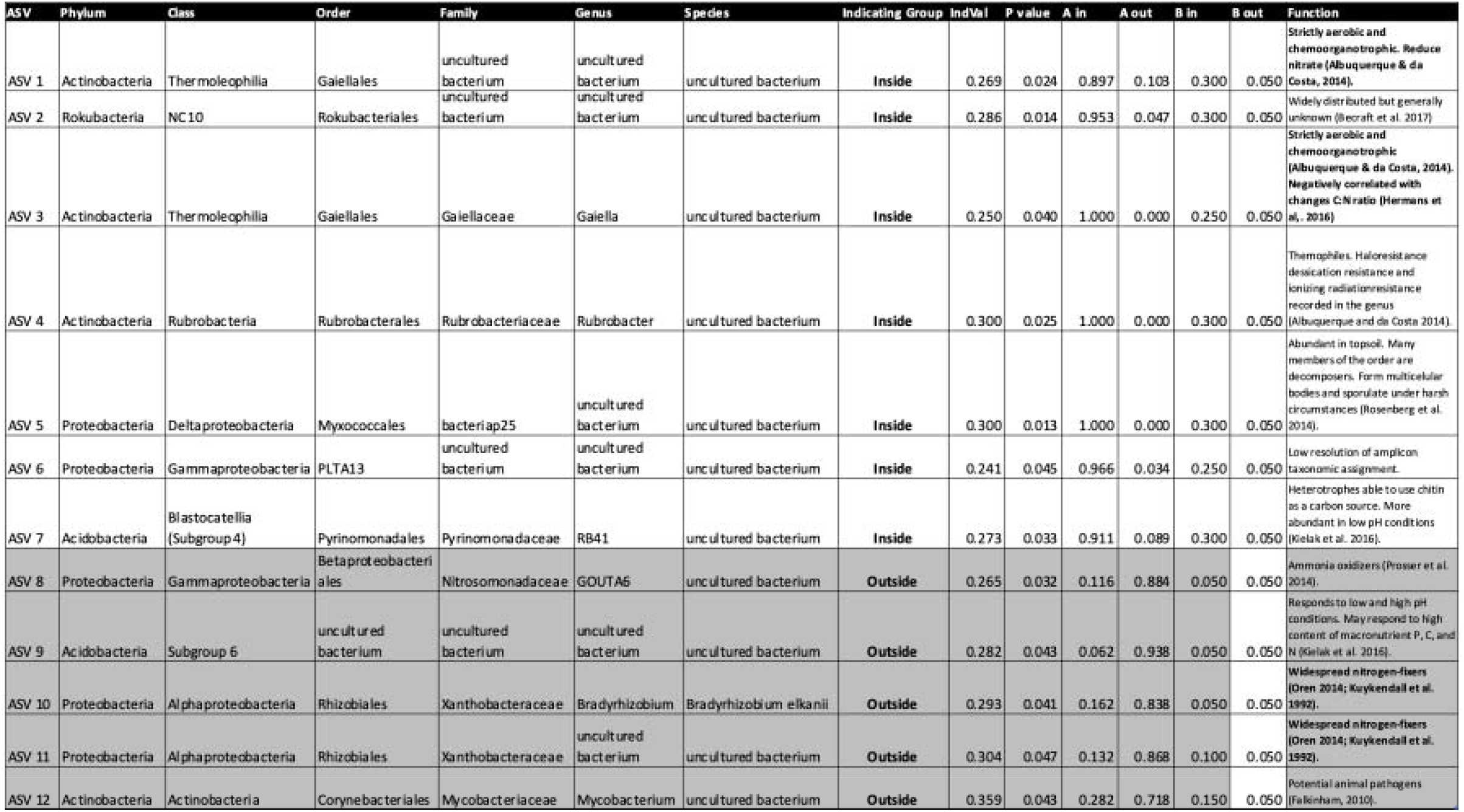
Indicator species associated with either invaded (In, white) or uninvaded (Out, Grey) areas. This table contains information about the phylum, class, order, family, genus, species, Indicating group (group that the species is associated with), IndVal (indicator value which quantifies the specificity of the relationship between the ASV and the associated group), P value, A in (the probability that the sample is from an invaded area given the fact that the species has been found in the sample aka the specificity), A out (the probability that the sample is from an uninvaded area given the fact that the species has been found in the sample aka the specificity), B in (the probability of finding the ASV in sites belonging to samples collected in invaded areas aka the fidelity), B out (the probability of finding the ASV in sites belonging to samples coinvllected in unvaded areas aka the fidelity).

### Root microbial communities

Roots of different host plant species varied significantly in AMF colonization (n = 226, Deviance _df: 5_= 18.233, P = 0.003). However, *A. petiolata* invasion (n = 226, Deviance _df: 1_= 3.7601, p-value > 0.05) and the interaction between host species and *A. petiolata* invasion did not predict AMF colonization (n = 226, Deviance _df: 5_= 3.760, p-value > 0.05) (**Fig. 6**). The incidence of root lesions also varied significantly among plant species (n = 226, Deviance _df: 5_ = 13.56, p-value < 0.05). *Alliaria petiolata* alone did not significantly explained differences in root lesions among plants (n = 226, Deviance _df: 1_ = 0.854, p-value > 0.05). However, there was a nearly significant interaction between *A. petiolata* and host plant species (n = 226, Deviance _df: 5_ = 9.705, p-value > 0.05), suggesting that *A. petiolata* invasion may have affected each host plant species differently. *Anemone americana, C. lutetiana* (Estimate: −0.066, CI = −0.260 to 0.128), *G. aparine* (Estimate = −0.080, CI = −0.265 to 0.105), *M. racemosum* (Estimate = 0.063, CI = -0.143 to 0.267), and *S. canadensis* (Estimate = 0.021, CI = −0.156 to 0.198) all had similar levels of lesions inside and outside *A. petiolata* invasion, but *G. robertianum* (Estimate = 1.091, CI = −0.618 to −0.052) had less lesions outside the invaded area than inside (**Fig. 7**).

**Figure 6.**
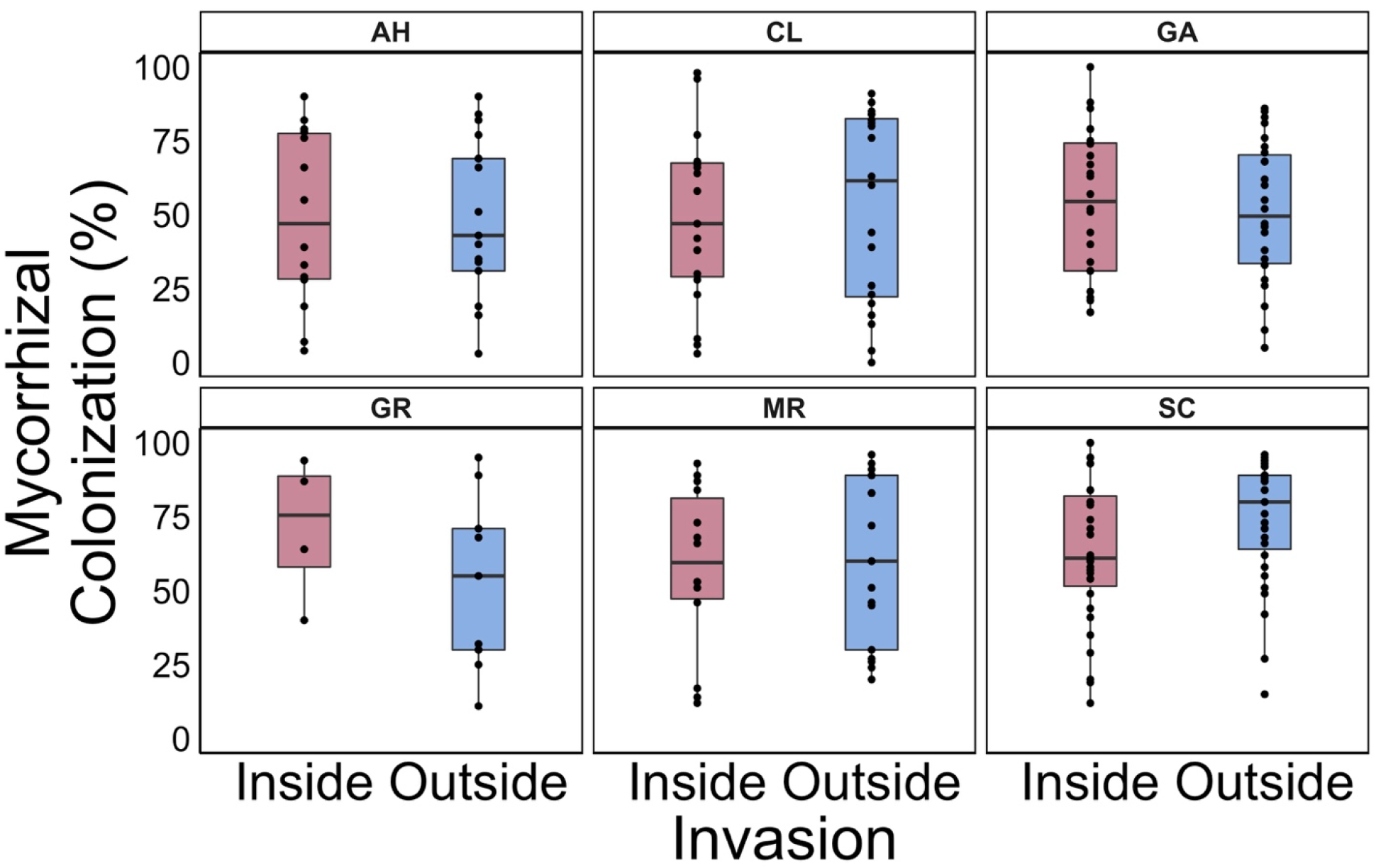
*Alliaria petiolata* invasion and total AMF colonization in co-occurring plant species. Plant species *Anemone americana* (AH), *Circaea lutetiana* (CL), *Galium aparine* (GA), *Geranium robertianum* (GR), *Maianthemum racemosum* (MR), and *Solidago canadensis* (SC) were collected from 10 populations of *A. petiolata* either inside (inside) or outside the invaded area (outside). Total colonization consists of the sum of all mycorrhizal structures at 100 root intersections per plant. The boxplots display the relationships between the presence of *A. petiolata* (Invasion) and AMF colonization (Mycorrhizal colonization). Arbuscular mycorrhizal fungal colonization was not different between invaded and uninvaded samples (P > 0.05) for any of the species sampled (P < 0.001).

**Figure 7.**
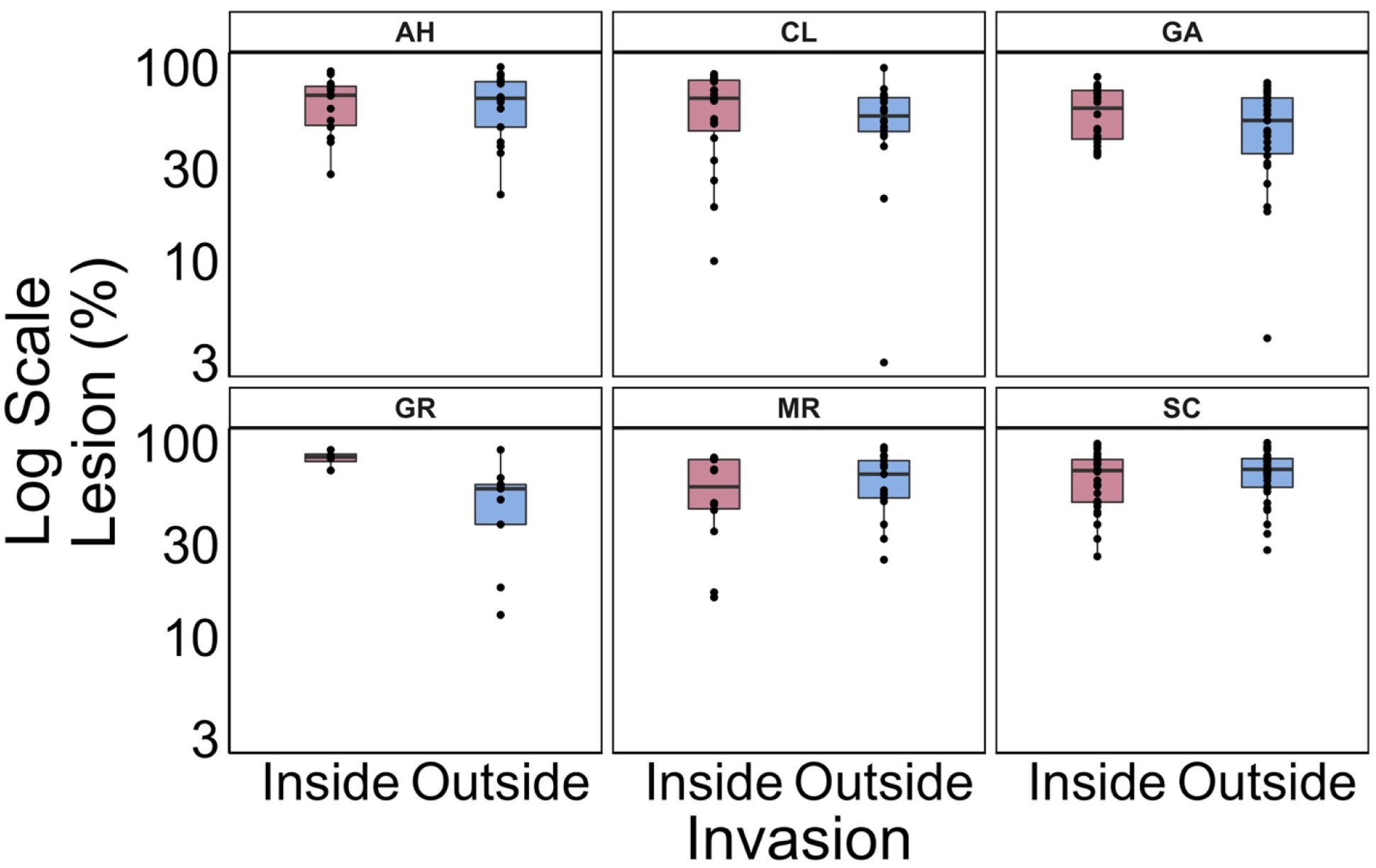
*Alliaria petiolata* invasion and lesions in roots of co-occurring plant species. Plant species *Anemone americana* (AH), *Circaea lutetiana* (CL), *Galium aparine* (GA), *Geranium robertianum* (GR), *Maianthemum racemosum* (MR), and *Solidago canadensis* (SC) were collected from 10 populations of *A. petiolata* either inside (inside) or outside the invaded area (outside). For each sample collected,weassessed the presence of structure or root damage related to pathogen presence at 100 random points on the roots to produce a measure of pathogen presence in percentage. The boxplots display the relationships between the presence of *A. petiolata* (Invasion) and the presence of pathogen related structures and root damage (Lesions). Note that the y axis is on the log scale to facilitate visualization. Root lesions were not evenly different between invaded and uninvaded samples (P > 0.05) but, *G. robertianum* had more lesions when collected inside garlic mustard (P < 0.001).

## Discussion

Our study is the first to examine the effect of *Alliaria petiolata* on root health and soil microbes under natural field conditions using molecular tools that are sensitive across a broad range of taxa. We confirmed that *A. petiolata* invasion is associated with shifts in soil microbial community composition but not as expected based on previously published laboratory studies. In previous studies, AMF suppression by biochemical compounds produced by *A. petiolata* has been implicated repeatedly as the key contributor to *A. petiolata*’s rapid invasion of Eastern North America (Barto et al., 2011; Callaway et al., 2008; Koch et al., 2011; Lankau, 2011a; Lankau, 2011b; Stinson et al., 2006). However, under natural field conditions we found that AMF community structure was unaffected by *A. petiolata* invasion. Conversely, we found EMF, saprotroph, nitrogen-fixing bacteria, and putative pathogen composition all shifted in the presence of this invasive species. Furthermore, we found marginally significant evidence for increased root lesions in *G. robertianum* co-occurring with *A. petiolata* invasion. These changes in functional composition may be important forces underlying the invasion of *A. petiolata* and its impact on ecosystem function.

We recorded that fungal richness increased significantly while bacterial richness remained constant in the presence of *A. petiolata*. Other studies on fungal communities have reported similar increases of overall fungal richness in the presence of *A. petiolata* and intermediate concentrations of its exudates (Anthony et al., 2017; Lankau, 2011a). Our results are also consistent with studies on bacteria reporting similar bacterial richness between *A. petiolata* invaded and uninvaded soils (Burke &Chan, 2010; Lankau, 2011a). Furthermore, we recorded a turnover in community composition of both soil bacteria and fungi in *A. petiolata* invaded soils when compared to uninvaded soils. Although we detected significant variation between technical replicates of 16S sequencing we included these replicates as part of our statistical models; previous studies have shown that variation between technical replicates are common and can be adequately accounted for in statistical models (Wen et al., 2017). Similarly, Burke and Chan (2010) recorded a difference in community composition of bacteria in soil near *A. petiolata* when compared to soil near native herbaceous species in late summer but did not measure fungal community composition. However, contrary to our findings, Lankau (2011a) noted no difference between bacterial and fungal species community composition of invaded and uninvaded soils as measured by TRFLP. The coarser data provided by TRFLP relative to direct sequencing may help explain the contradicting results found in the literature. In contrast with previous studies of the microbial community associated with *A. petiolata* invasion (Anthony et al., 2017; Barto et al., 2011; Burke, 2008; Burke & Chan 2010) we sequenced the V4 region of the 16S ribosomal RNA gene to detect bacteria and archaea (Walters et al., 2016) and part of the small subunit ribosomal RNA encoding gene, the internal transcribed spacer DNA, and part of the large-subunit ribosomal RNA region of the genome to detect fungi (Gao et al., 2019; Taylor et al., 2016). Our approach, which provides a more detailed representation of the microbial community compared to previous studies using TRFLP, enable the detection of spatial turnover of bacterial species composition and an increase in richness with *A. petiolata* invasion. We were also able to assign potential functional attributes based on phylogenetic assignment to infer changes to ecosystem processes such as nutrient cycling.

Due to the observational nature of our experiment, we cannot determine with certainty whether *A. petiolata* is invading the ecosystem as a passenger or driver of soil microbial change, a question fittingly named the Passenger-Driver hypothesis (Bauer, 2012; MacDougall & Turkington, 2005). However, our data are consistent with the existence of potential causal links between the removal of *A. petiolata* and changes in microbial communities (Burke et al. 2019; Lankau, Bauer, Anderson, & Anderson, 2014) as well as processes such as litter decomposition (Rodgers et al., 2008), lending support to the ‘Driver’ hypothesis in invasion biology. Additionally, the proximity and similarity in soil properties between our invaded and uninvaded samples suggests that the differences in microbial communities are likely a result from the dispersal limitations of *A. petiolata* rather than the inability of its microbiome to colonize adjacent soils.

Despite the change in overall fungal community composition accompanying the presence of *A. petiolata*, we found no evidence for the suppressive effect of *A. petiolata* on AMF diversity or AMF root colonization. These data are comparable to those of other studies on naturally occurring populations of *A. petiolata* providing poor support for changes in AMF community composition (Barto et al., 2011; Burke, 2008) but contrast with studies that have detected a reduction in soil AMF richness (Lankau, 2011b). Part of the difference between our results and that of Barto et al (2011) could be attributed to the different canopy cover of the two sites. Our study site is dominated by tree species that are both AMF- and EMF-dependent as opposed to the AMF-dominated species found in the site studied by Barto et al. (2011). The difference between the mycorrhizal colonization of the sites could result in a stronger AMF response in the AMF-dominated sites. Nonetheless, while leaf and root extracts of *A. petiolata* have been shown to inhibit AMF growth (Koch et al., 2011; Roberts & Anderson, 2001; Stinson et al., 2006) and could reduce AMF colonization in *A. petiolata* invaded soils, our data raises questions about the ecological significance of AMF suppression as a putative mechanism of invasion. Our study reinforces the importance of conducting observational field studies using molecular tools aimed at detecting ecologically significant microbial functional groups.

The inconsistent effect of *A. petiolata* on AMF communities may be attributed to a breakdown of secondary metabolites by edaphic processes. Barto & Cipollini (2009) examined biochemicals implicated in *A. petiolata* mycorrhizal suppression from bulk soils collected in the field but found that glucosinolates were undetectable under natural field conditions. However, some potentially bioactive flavonoid derivatives could be detected a few times during the season. Still, both flavonoids and glucosinolates had half-lives of merely a few hours in field collected soils. As such, the effects of *A. petiolata* on AMF communities could be dampened by natural processes such as microbial degradation or seasonal fluctuations of biochemical release. Alternatively, intraspecific variation in glucosinolate production could explain the variability in the effect of *A. petiolata* on AMF reported in the literature. Lankau (2011a) reported that the concentration of glucosinolates in *A. petiolata* increased with population age but decreased in populations established for more than 60 years. The populations we studied are relatively young (less than 20 years) and vary in density; thus shifts in AMF may simply not be detectable yet. Nonetheless, the reoccurrence of field results that do not support the existence of detrimental effects of *A. petiolata* on AMF communities, especially in newly invaded areas, should bring into question the claim that AMF suppression is one of the primary mechanisms of invasion by this species. We posit that to understand how *A. petiolata* is changing ecosystems, the focus of research must move away from an *A. petiolata*-AMF-native plant relationship to include other functional groups.

We detected changes in ectomycorrhizal community composition in *A. petiolata* invaded sites. While we cannot rule out the possibility of direct *A. petiolata* mediated effects, these changes could result indirectly from shifts in decomposition rates associated with *A. petiolata* invasion (Rodgers et al., 2008). *Alliaria petiolata* leaf litter is higher in nitrogen than the leaf litter found in uninvaded forests (Rodgers et al., 2008). Ectomycorrhizal fungi have the potential to decompose soil organic matter, targeting the release of nitrogen (Lindahl & Tunlid, 2014; Talbot et al., 2013). In fact, EMF communities are known to respond to changes in leaf litter quality and composition (Conn & Dighton, 2000; Eisenhauer, Reich & Isbell, 2012). However, EMF have evolved independently several times from saprotrophic ancestors (Hibbett, Gilbert & Donoghue, 2000). The genetic potential of EMF to decompose soil organic matter and release nitrogen varies widely by lineage (Pellitier & Zak, 2017). Therefore, the changes in nitrogen availability and decomposition brought about by *A. petiolata* invasion are unlikely to affect every EMF species in the same way. This hypothesis is consistent with our observations that EMF communities were not suppressed by *A. petiolata*, but rather experienced a change in composition, which is consistent with the species-specific response of EMF to *A. petiolata* invasion recorded in other studies (Castellano & Gorchov, 2012; Anthony et al., 2017). In addition, past observations by Wolfe et al. (2008) support the hypothesis that changes in decomposition and soil nitrogen availability due to *A. petiolata* leaf litter, rather than *A. petiolata* root exudates, impact EMF community composition. They found that *A. petiolata* invasion was associated with a reduced abundance of EMF colonizing root tips in forest populations but recorded no such difference between soil conditioning by growth of *A. petiolata* as compared to conditioning by the native *Impatiens capensis* under greenhouse conditions. One of the major differences between greenhouse and field studies is that under natural conditions the EMF community can be indirectly affected by the decomposition of *A. petiolata* leaf litter but in the greenhouse only direct effects of *A. petiolata* exudates have been considered since the leaf litter was never allowed to decompose. Therefore, the shifts in EMF community observed here are likely to result from an indirect consequence of *A. petiolata* invasion rather than a direct effect as previously suggested. The potential threat of *A. petiolata* invasion to EMF-dependent plant species makes this an important avenue to explore for future research.

Invasive plant species influence soil nutrient cycling and decomposition (Ashton, Hyatt, Howe, Gurevitch & Lerdau, 2005; Ehrenfeld, 2003), which in turn can affect soil decomposers. Our results show that the community composition of fungal decomposers was altered in presence of *A. petiolata*. We did not measure decomposition rates directly but rather recorded an increase in richness of saprotrophs. This is consistent with the results reported by Rodgers et al. (2008) where litter bags of *A. petiolata* first-year rosette leaves placed in uninvaded forest areas decomposed, released nitrogen, and stimulated the decomposition of leaf litter from dominant native trees faster than leaf litter bags of the native *Acer saccharum* and *Impatiens capensis*. Rodgers et al. (2008) observed an increase in nitrogen immobilization but still higher nitrogen availability in invaded than in uninvaded soil. Therefore, based on supporting literature and the results found in this study, *A. petiolata* appears to be an important driver of changes in biogeochemical cycles of the forest understory.

Changes in microbial communities associated with nutrient cycling in *A. petiolata* invaded soils were not limited to fungi. We found that the majority of the bacterial species that differed significantly between invaded and uninvaded soils are associated with the nitrogen cycle. Of the five species identified as statistically significant indicators of invaded soil, two were symbiotic nitrogen-fixing bacteria and one was an ammonia oxidizer. There was a significant decrease in the relative abundance of nitrogen-fixing bacteria species co-occurring with *A. petiolata*, suggesting that these species, and their plant hosts, responded negatively to *A. petiolata*. In laboratory experiments, Portales-Reyes, Van Doornik, Schultheis, & Suwa (2015) found that chemical compounds associated with *A. petiolata* affected associations with rhizobia negatively. Conversely, because *A. petiolata* invasion increases nitrogen availability, the suppression of some Xanthobacteraceae could be a result of the *A. petiolata* leaf litter decomposition rather than allelopathic inhibitions; an increase in nitrogen availability caused a reduction in rhizobia nodulation (Lu et al., 2011; Streeter & Wong, 1998). Although the exact mechanism suppressing the nitrogen-fixing symbiont is unknown, microbes involved in the nitrogen cycle were clearly affected by *A. petiolata*. Nitrogen is a critical limiting nutrient for plant growth (Vitousek & Howarth, 1991), which means that it can modulate net primary production and shape plant community composition (LeBauer & Treseder, 2008). Therefore, the effect of *A. petiolata* on soil nitrogen cycling is of particular interest when considering long term changes to terrestrial communities and should be investigated further.

The accumulation of pathogen in non-native species and the subsequent spillback onto native species has been proposed as a mechanism of invasion (Mangla & Callaway, 2008; Flory & Clay, 2013). In accordance with Anthony et al. (2017), we recorded an increase in putative fungal plant pathogens in soil collected within *A. petiolata* populations, suggesting that soil around the invasive plant is accumulating pathogens. However, the long-term accumulation of pathogens in soils colonized by invasive plants can lead to either the decline of invasive species or the decline of its native neighbors through spillover effects (Flory & Clay, 2013). The family Spizellomycetaceae (dominated by *Spizellomyces palustris* and *S. lactosolyticus*), Mortierellaceae (all from the genus *Mortierella*), and Nectriaceae (mainly *Fusarium solani*) dominated the pathogen guild in *A. petiolata* invaded soils, but there is a lack of information on the ecology and host preference of these fungal pathogens. Therefore, we cannot draw direct conclusions from sequence data on whether the increase in these putative pathogens could be detrimental to the invasive plant or its native neighbors. Instead we found some evidence that *A. petiolata* may reduce the root health of some cohabitating plant species as detected by quantification of root lesions. This suggests that pathogen spillover may play a role in invasion which should be investigated further. Heterotrophic fungi can overlap in their physiological capacity for pathogenic and saprotrophic strategy (Newton, Fitt, Atkins, Walters & Daniell, 2010; Olson et al., 2012). Specifically, the family Mortierellaceae dominated the saprotroph community and was also the third most abundant family of pathogens in *A. petiolata* invaded soils. As such, the accumulation of saprotrophs that are facultative pathogens could be related to or exacerbate the accumulation of pathogens associated with *A. petiolata* invasion. Thus, *A. petiolata* seems to be promoting the proliferation of potentially facultative pathogens that may spread to co-occurring native plants and reduce their health and fitness.

In conclusion, our study represents the first comprehensive census of microbial community changes related to *A. petiolata* invasion in a natural setting. We found little evidence to support the hypothesis that AMF suppression is a key driver of *A. petiolata* invasion. Instead, we found evidence that invasive plants have the potential to act as driver of invasion through changes in the soil microbiome. The proliferation of pathogens, changes in ectomycorrhizal community composition, and changes in microbial groups involved in nitrogen cycling must be further examined to increase our understanding of the impact of *A. petiolata* invasion on ecosystems.

## Supporting information

Supplementary Material about microbial families 16S and ITS

Supplementary Material about microbial functional group assignments ITS

Supplementary Material about microbial phylum 16S and ITS

## Acknowledgments

This research was supported by grants from the Natural Sciences and Engineering Research Council of Canada to Pedro M. Antunes and Robert I. Colautti. We thank Almira Siew for assistance with the laboratory work. We are also grateful to Dr. Virginia Walker, Dr. Paul Grogan, and Dr. William Plaxton from Queen’s University for comments that greatly improved the manuscript.

## Data Accessibility

The DNA sequences that support the findings of this study are openly available in Sequence Read Archive (SRA) database accession number PRJNA551217. [Reviewer link: https://dataview.ncbi.nlm.nih.gov/object/PRJNA551217?reviewer=b5h0i6ndv1a93gs0cois0h4moj]. The Root lesion and mycorrhiza data, the bioinformatic pipeline, and the reproducible analysis (R code) that support the findings of this study are openly available in Dryad at http://doi.org/10.5061/dryad.hqbzkh1bk.

## Author’s contributions

KD designed and performed the experiments and analyzed the data with support from RC and PA. AG assisted with root lesion and mycorrhiza measurements. KD wrote the manuscript in consultation RC, PA, and AG. Financial support came from RC and PA.

## Notes

### Competing Interest Statement

The authors have declared no competing interest.

