## Supplementary Material for "Functional shifts of soil microbial communities associated with *Alliaria petiolata* invasion"

**Affiliations:**

**Table S.1** Latitudinal and longitudinal coordinates of the plot location

| Population Number | Latitude | Longitude |
| --- | --- | --- |
| 1 | N44° 34.511' | W76° 19.788' |
| 2 | N44° 34.060' | W76° 19.530' |
| 3 | N44° 33.951' | W76° 19.489' |
| 5 | N44° 33.100' | W76° 21.433' |
| 7 | N44° 33.069' | W76° 21.691' |
| 8 | N44° 33.204' | W76° 21.943' |
| 9 | N44° 33.050' | W76° 22.241' |
| 10 | N44° 32.840' | W76° 21.982' |
| 13 | N44° 35.052' | W76° 22.615' |
| 14 | N44° 34.772' | W76° 22.511' |

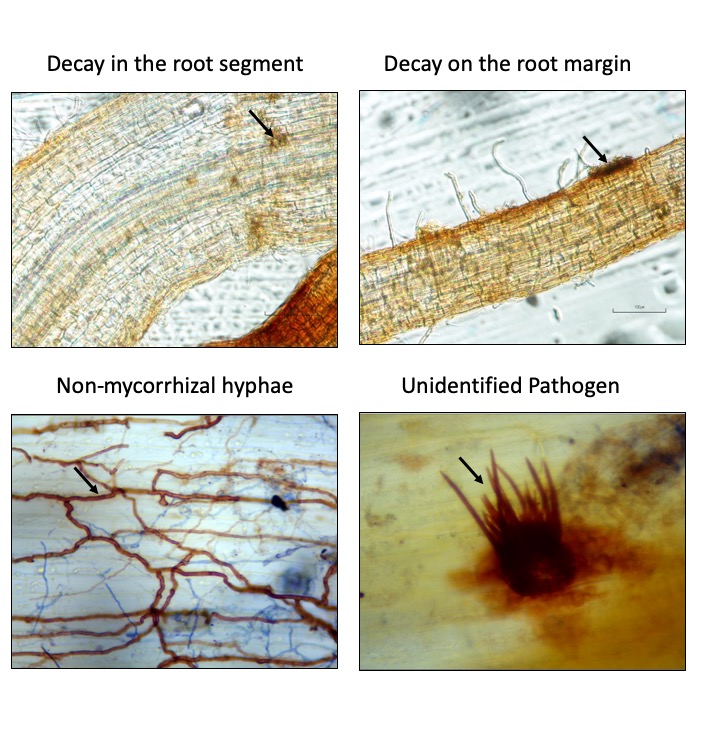

**Figure S1.** Reference images of signs of pathogen infection (referred to as lesions in the text). **A)** A black arrow points to an area of decay in the root segment. **B)** A black arrow points to an area of decay on the root edge. Decay inside and on the edge of the root were counted as one form of decay but are represented separately here for reference. **C)** A black arrow points to the brown and septate non-mycorrhizal hyphae. **D)** A black arrow points to an unidentified pathogen.

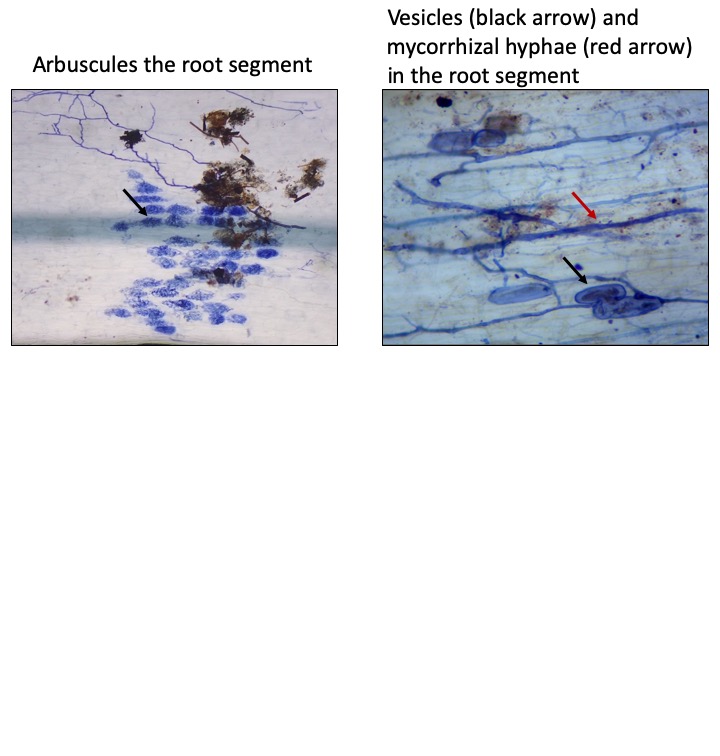

**Figure S2.** Reference images of mycorrhizal colonization. **A)** A black arrow points towards an arbuscule, dyed blue. **B)** A black arrow points to a vesicle in blue, and a red arrow points to a non-septate mycorrhizal hypha. Hyphae and vesicles were counted as separate structures but are on the same image here for reference.

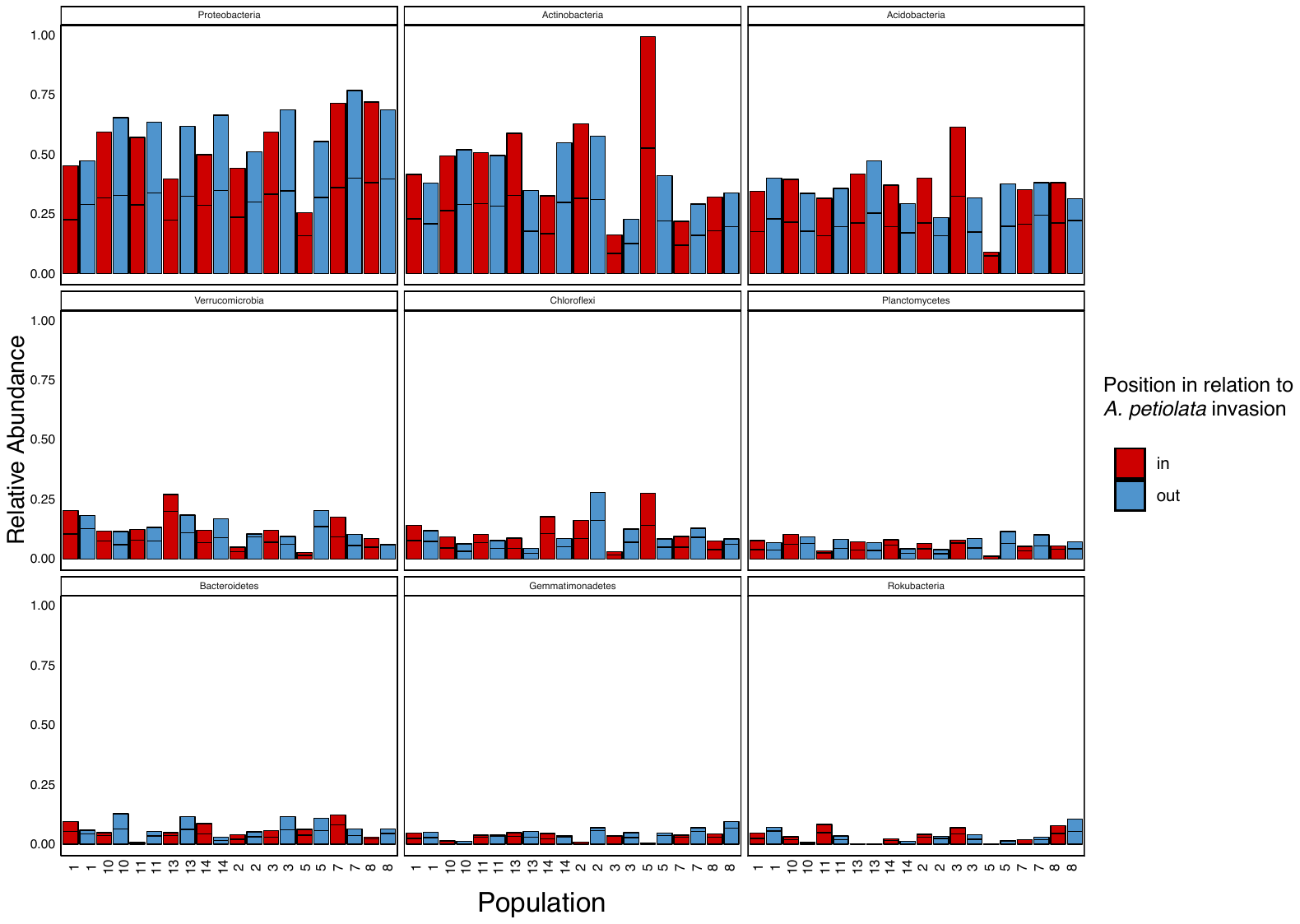

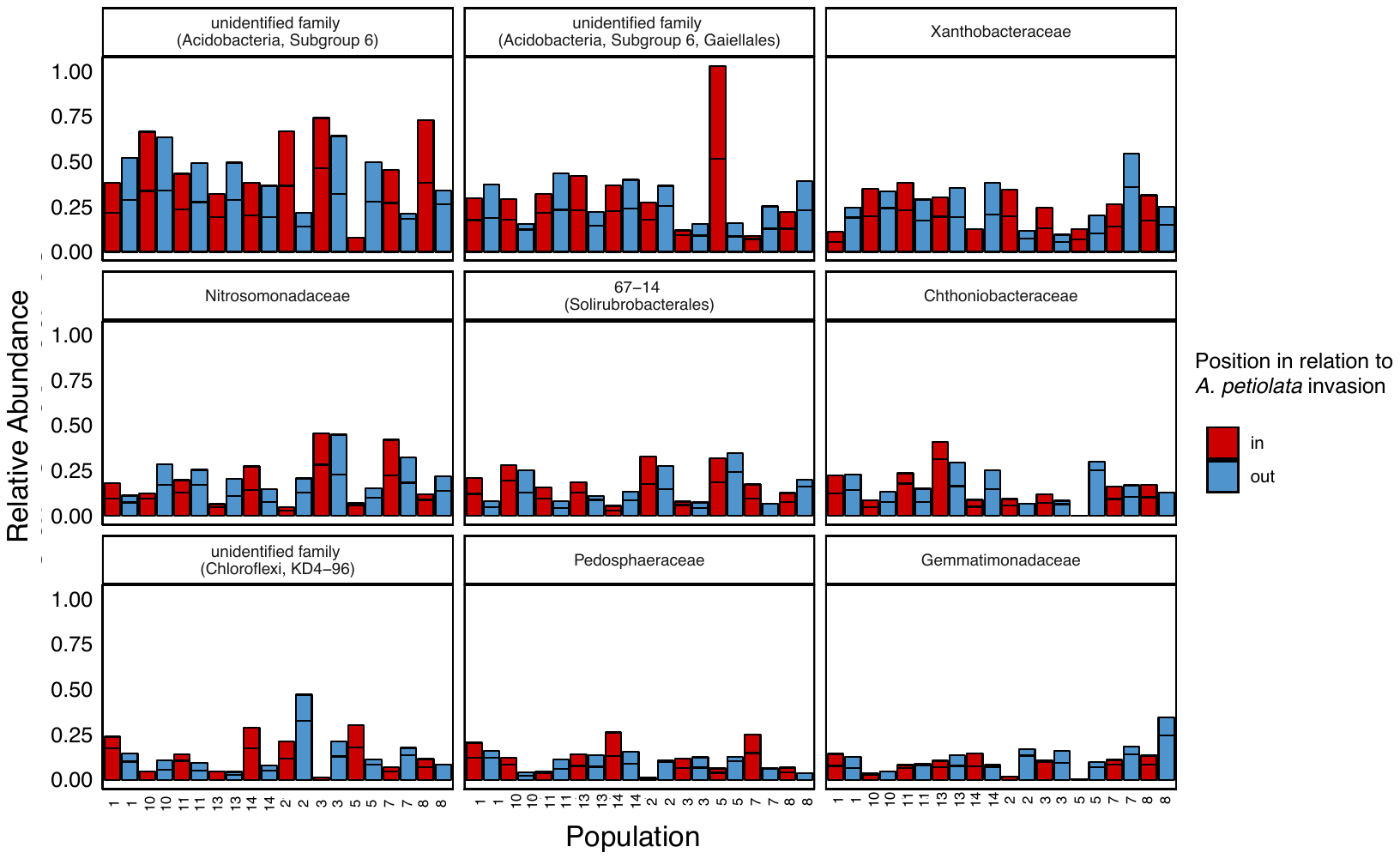

**Figure S3. A)** Each facet represents one of the nine phyla of bacteria that were most common in all the samples, from most to least common. **B)** Each facet represents one of the nine families of bacteria that were most common in all the samples, from most to least common. Samples from the same population (numbered) are next to each other. For both A) and B), in each population, one sample was collected inside (in, red) and one outside, but near, *A. petiolata* (out, blue). The bars show the relative abundance of the phylum in each sample, with technical replicates stacked but separated by black lines.

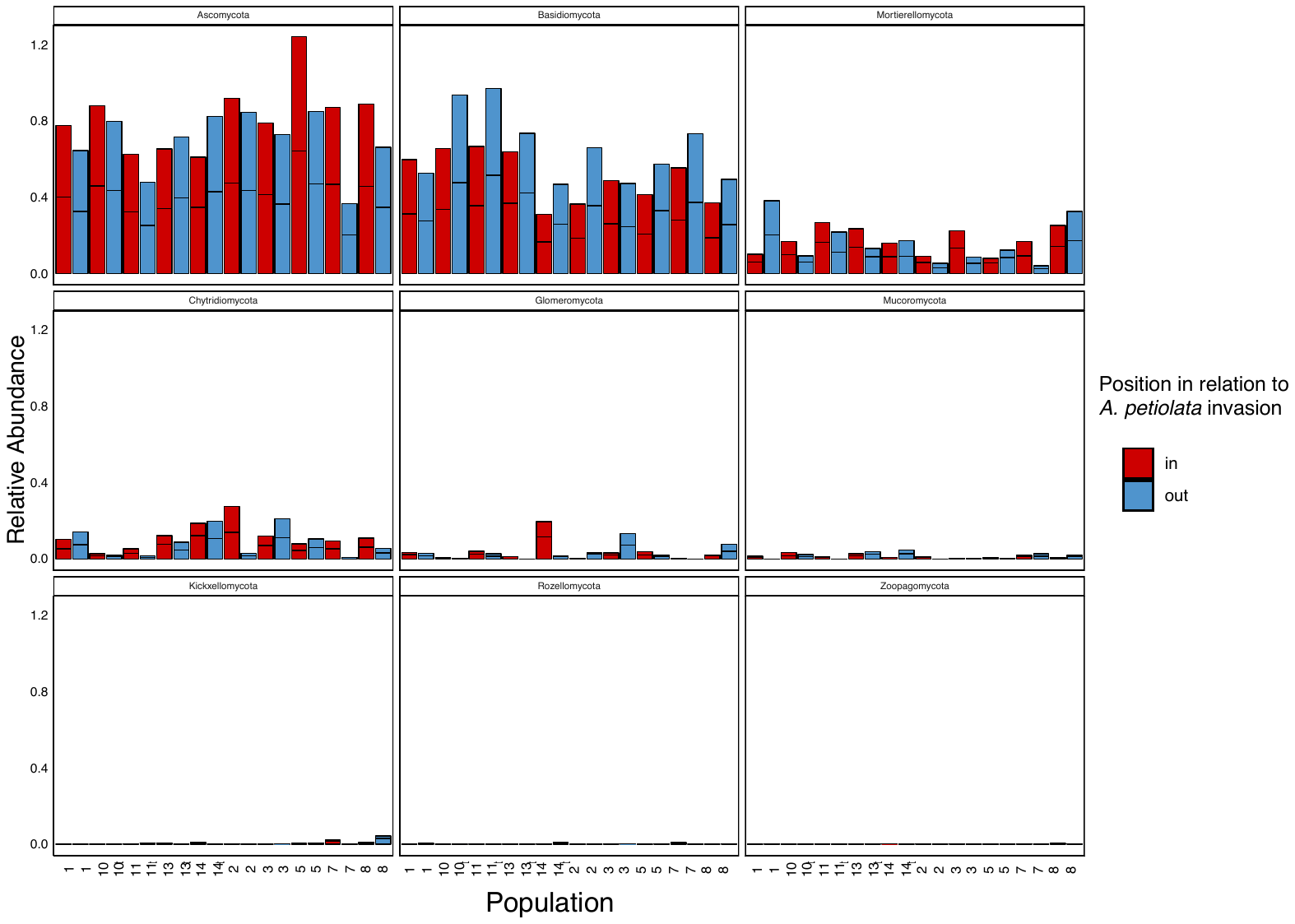

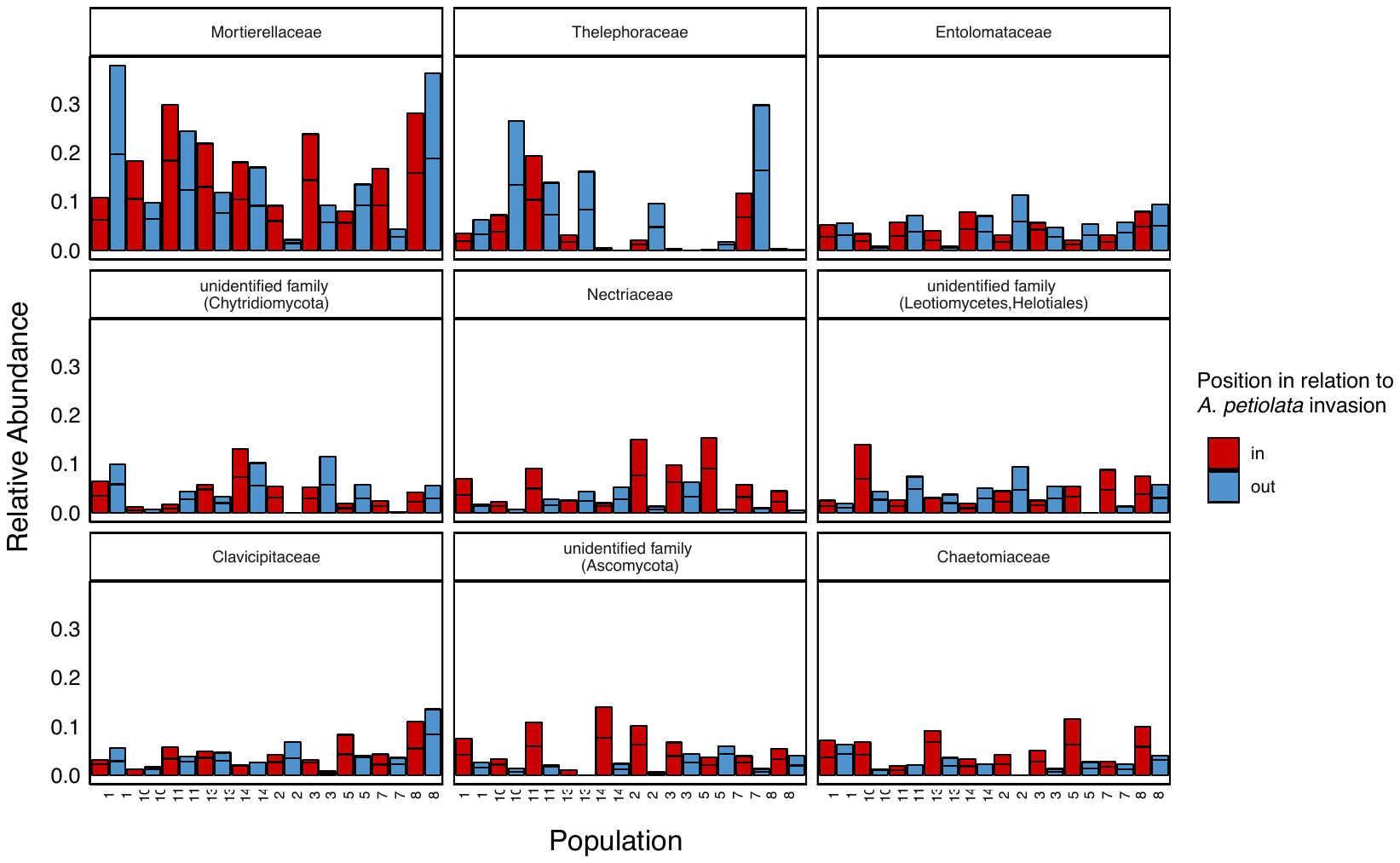

**Figure S4. A)** Each facet represents one of the nine phyla of fungi that were most common in all the samples, from most to least common. **B)** Each facet represents one of the nine families of fungi that were most common in all the samples, from most to least common. Samples from the same population (numbered) are next to each other. For both A) and B), in each population, one sample was collected inside (in, red) and one outside, but near, *A. petiolata* (out, blue). The bars show the relative abundance of the phylum in each sample, with technical replicates stacked but separated by black lines.

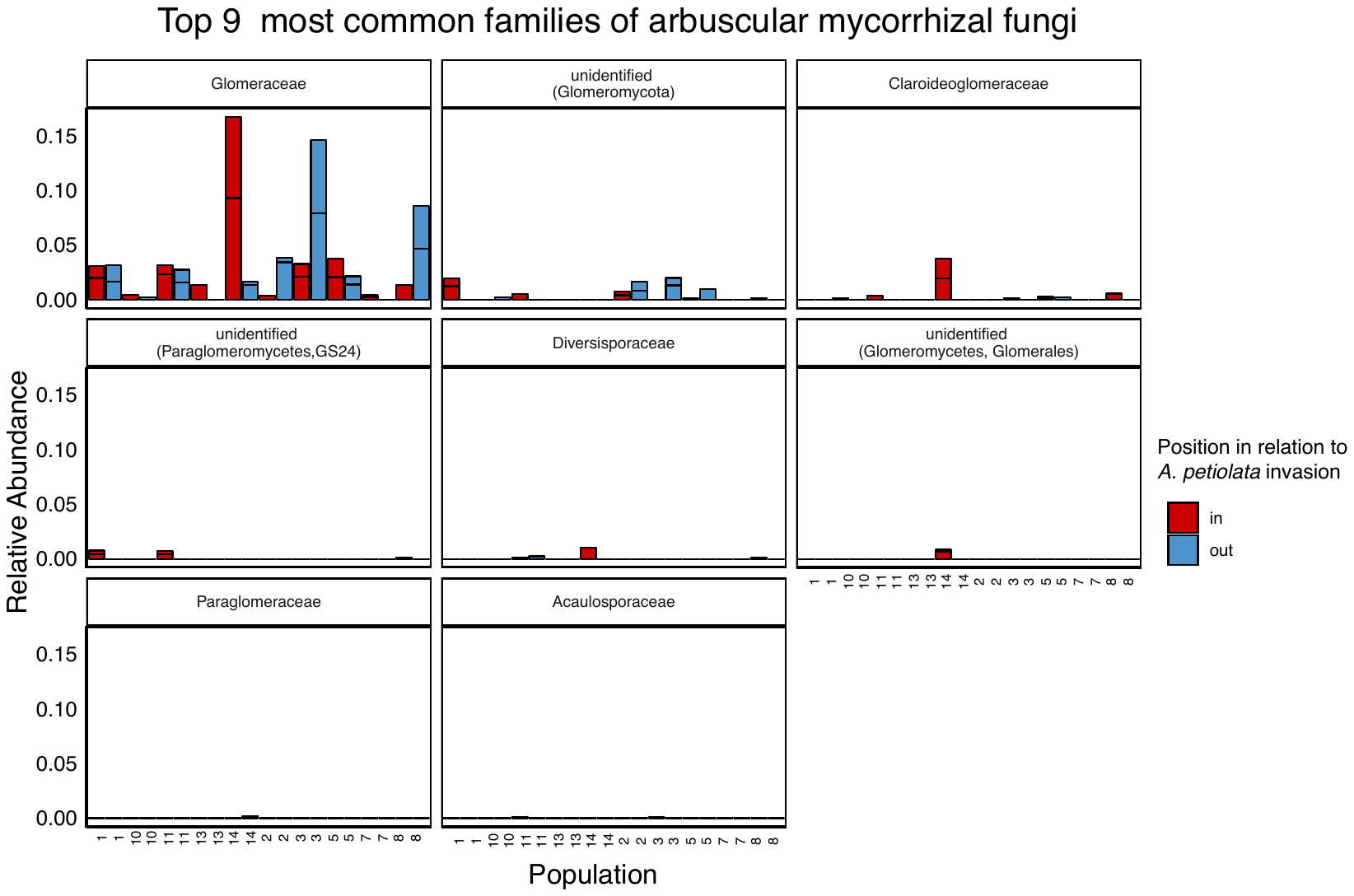

**Figure S5.** Each facet represents one of the eight families of arbuscular mycorrhizal fungi that were most common in all the samples, from most to least common. Samples from the same population (numbered) are next to each other. In each population, one sample was collected inside (in, red) and one outside, but near, *A. petiolata* (out, blue). The bars show the relative abundance of the phylum in each sample, with technical replicates stacked but separated by black lines..

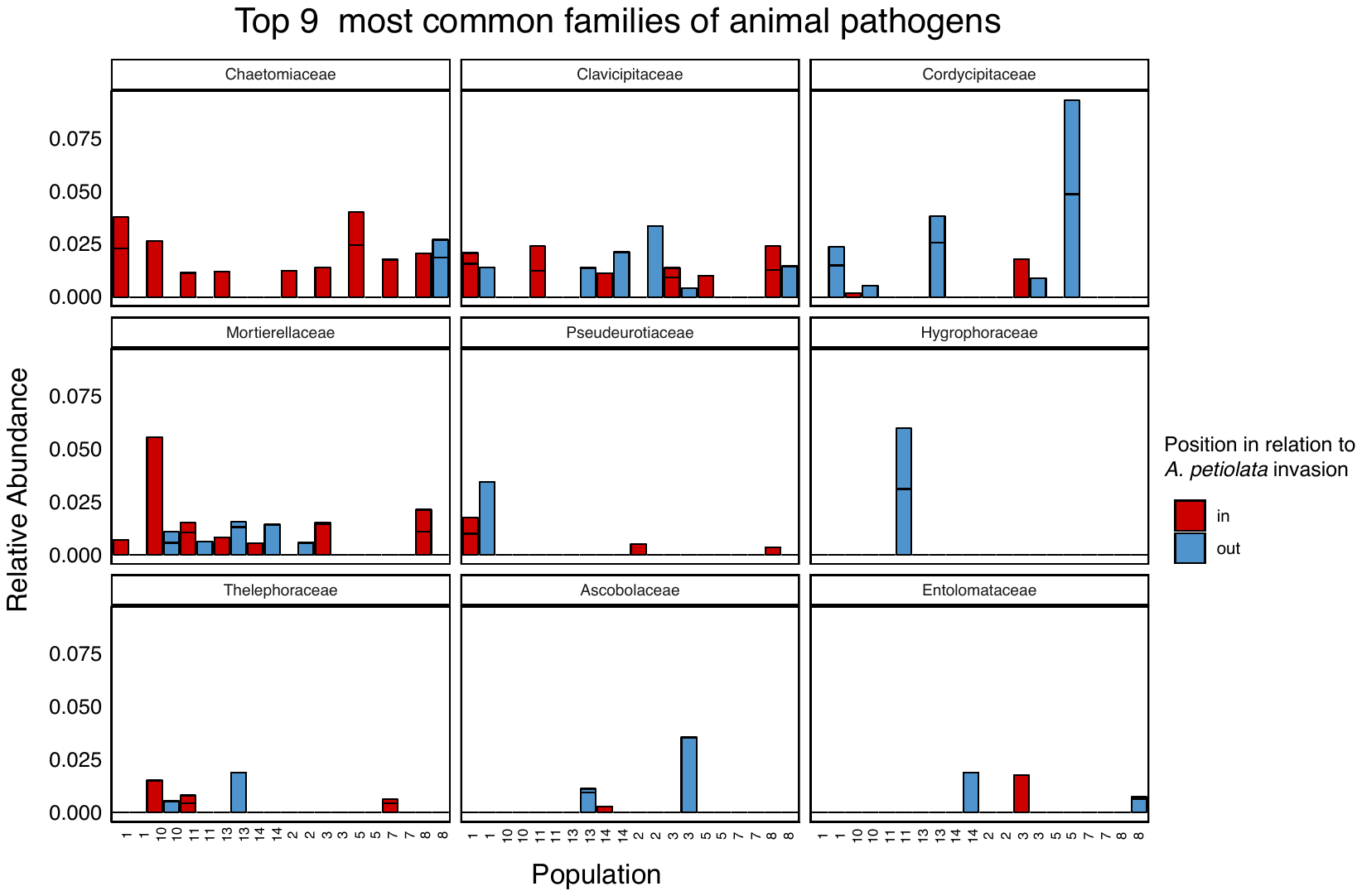

**Figure S6.** Each facet represents one of the nine families classified as animal pathogens that were most common in all the samples, from most to least common. Samples from the same population (numbered) are next to each other. In each population, one sample was collected inside (in, red) and one outside, but near, *A. petiolata* (out, blue). The bars show the relative abundance of the phylum in each sample, with technical replicates stacked but separated by black lines.

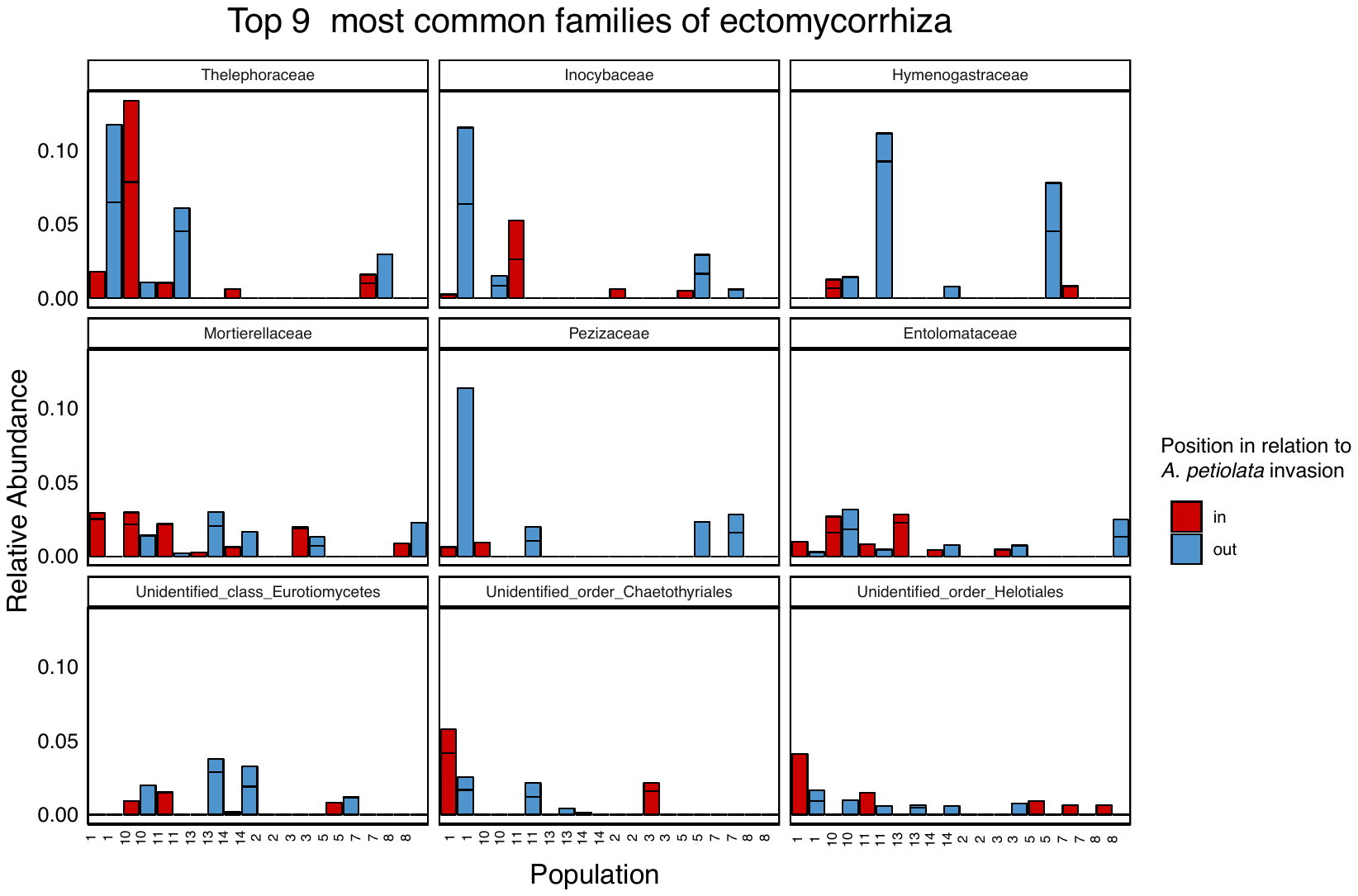

**Figure S7.** Each facet represents one of the nine families classified as ectomycorrhiza that were most common in all the samples, from most to least common. Samples from the same population (numbered) are next to each other. In each population, one sample was collected inside (in, red) and one outside, but near, *A. petiolata* (out, blue). The bars show the relative abundance of the phylum in each sample, with technical replicates stacked but separated by black lines.

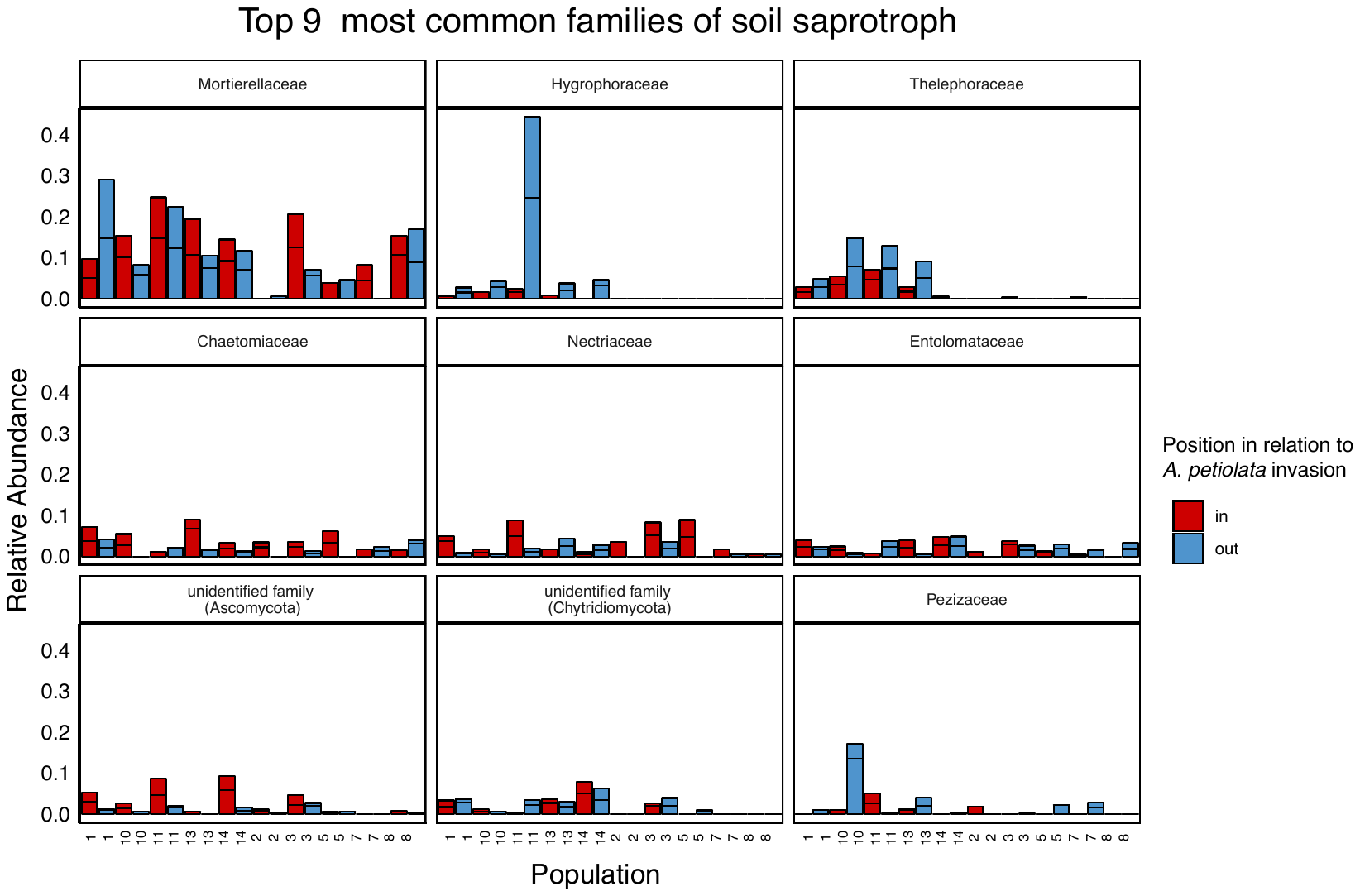

**Figure S8.** Each facet represents one of the nine families classified as soil saprotrophs that were most common in all the samples, from most to least common. Samples from the same population (numbered) are next to each other. In each population, one sample was collected inside (in, red) and one outside, but near, *A. petiolata* (out, blue). The bars show the relative abundance of the phylum in each sample, with technical replicates stacked but separated by black lines.

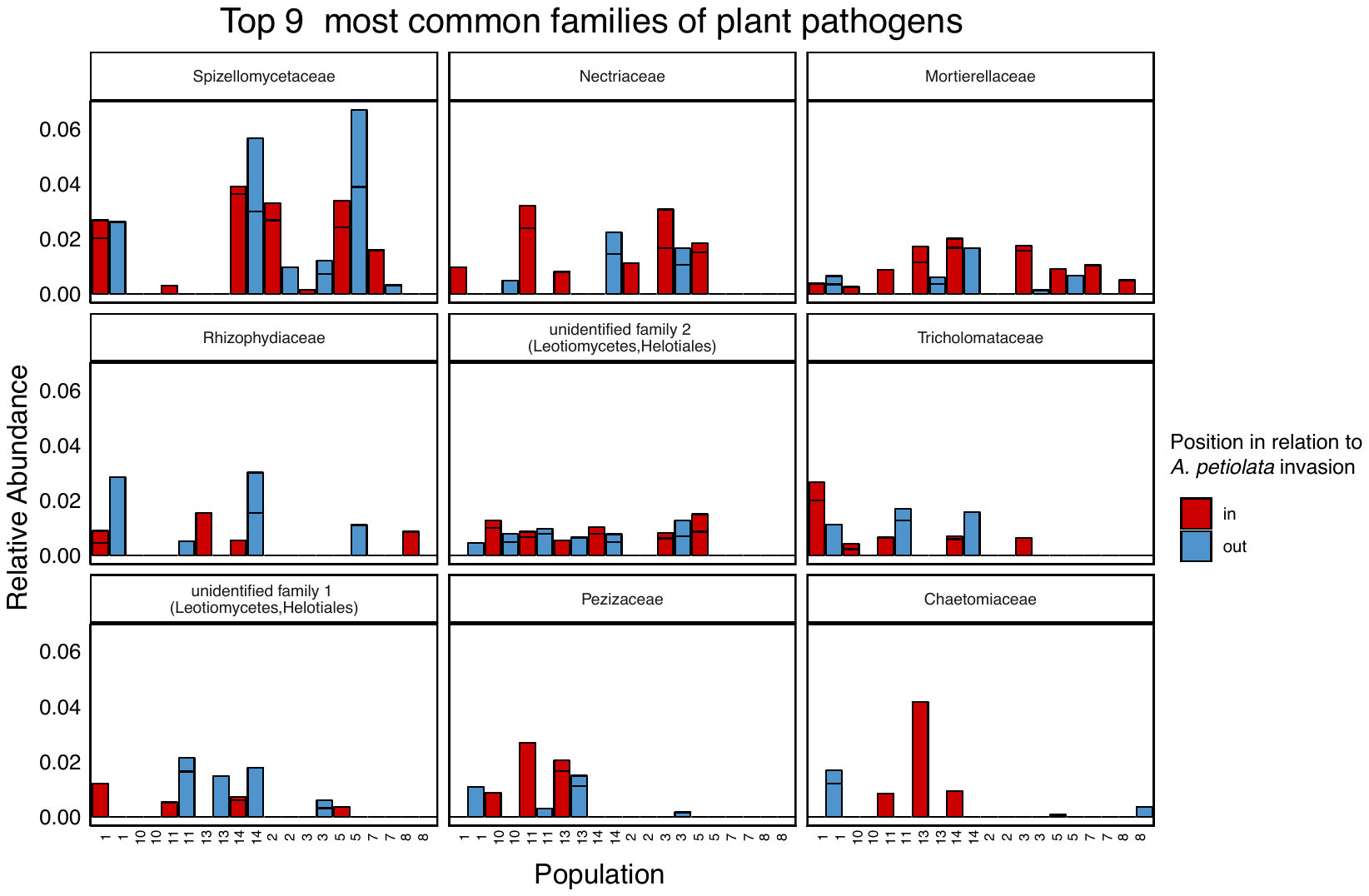

**Figure S9.** Each facet represents one of the nine families classified as plant pathogens that were most common in all the samples, from most to least common. Samples from the same population (numbered) are next to each other. In each population, one sample was collected inside (in, red) and one outside, but near, *A. petiolata* (out, blue). The bars show the relative abundance of the phylum in each sample, with technical replicates stacked but separated by black lines.

A)

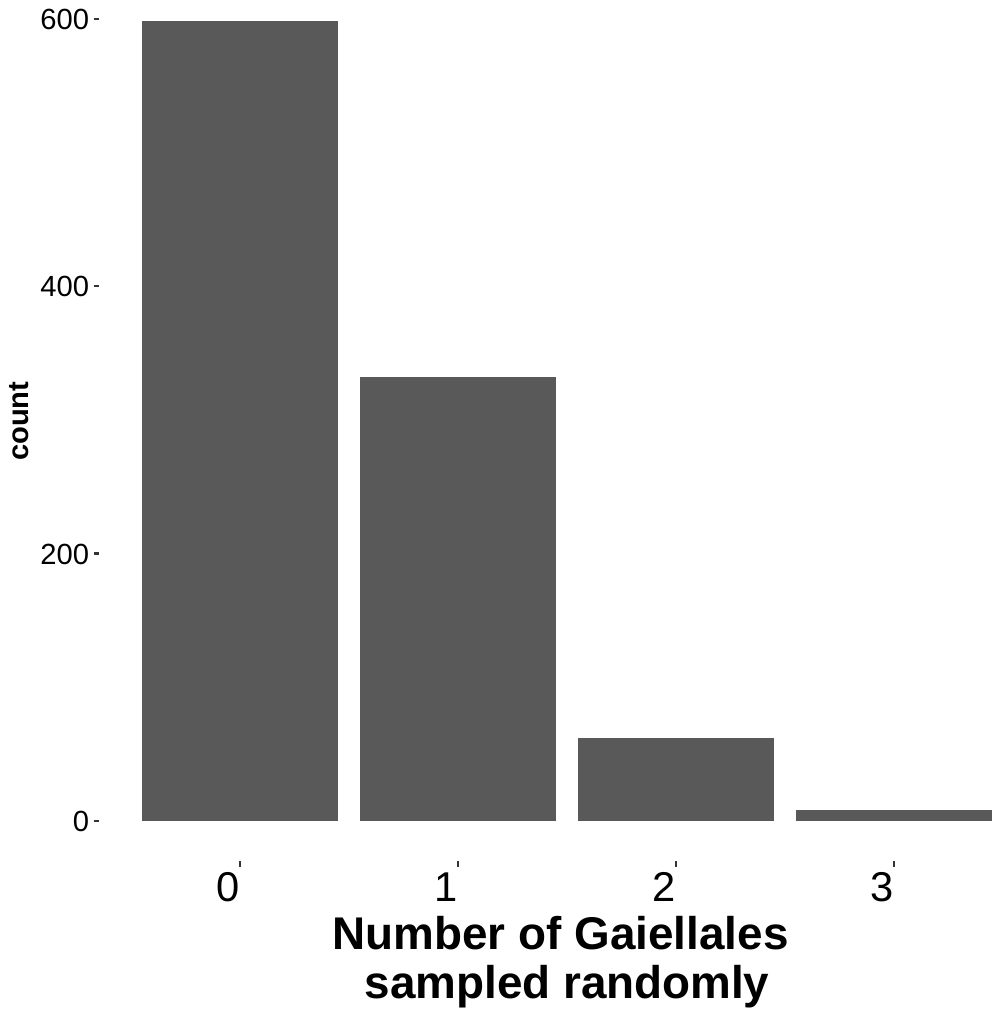

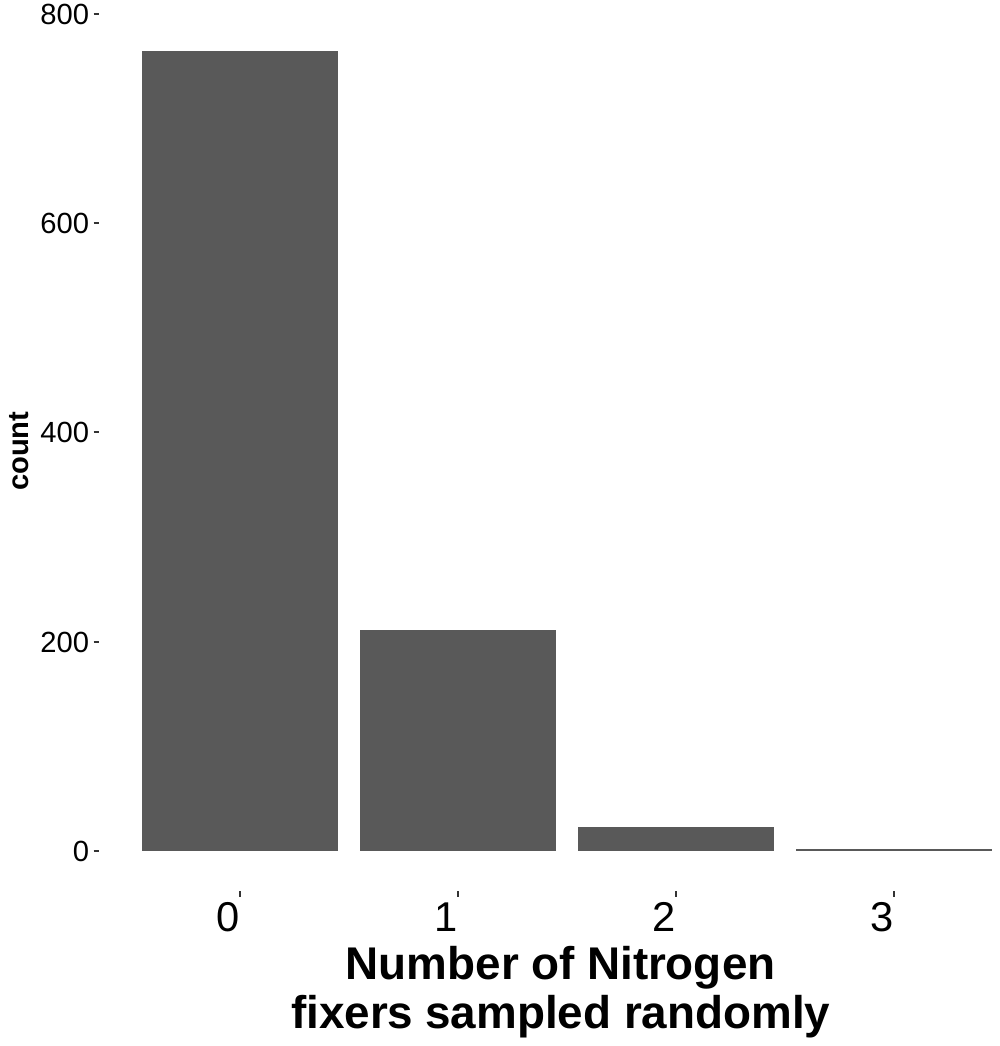

B)

**Figure S10. A)** The null distributions for the simulation of Gaiellales in *A. petiolata* invaded soil. The null distribution was produced by randomly sampling 7 ASVs for 1000 iterations and recording for each iteration how many of the sampled ASVs were Gaiellales. The x-axis represents the number of Gaiellales randomly sampled and the y-axis represents how many times this number of Gaiellales occurred in the 1000 iterations. **B)** Null distributions for the suppression of Nitrogen-Fixers in *A. petiolata* invaded soil. The null distribution was produced by randomly sampling 5 ASVs for 1000 iterations and recording for each iteration how many of the sampled ASVs were Nitrogen-Fixers. The x-axis represents the number of Nitrogen-Fixers randomly sampled and the y-axis represents how many times this number of Nitrogen-Fixers occurred in the 1000 iterations.

**Protocol for the measurement of soil pH:**

1. Sieve soil through 4mm sieve to remove rocks, roots, etc.

2. Add 10g of soil from each sample into a 50 mL falcon tube.

3. Add 20 mL of deionized water, for a 2:1 water to soil ratio. Vortex for 15

seconds.

4. Shake or vortex every fifteen minutes for an hour.

5. Shake before placing pH electrode into the soil solution. Wait for reading

to stabilize.

6. Repeat with all samples, making sure to clean the electrode between

samples.

**Protocol for the measurement of soil texture with a hydrometer**

1. Weigh 40 g of air-dried, fine texture soil (100 g if the soil is loamy sand) into a dispersion cup.
2. Add deionized (DI) water to make up to 300 mL.

*Note: Use DI water that is at room temperature*

3. Add 50 mL of sodium hexametaphosphate solution (100 mL if the sample weight is 100 g) to the dispersion blender cup.
4. Attach the dispersion cup to the electric mixer.
5. Mix the suspension for 5 minutes, taking care not to hit the bottom of the cup with the mixing rod.
6. Transfer the soil suspension quantitatively into a sedimentation cylinder.
7. Rinse the dispersion cup with DI water and add the rinse to the cylinder.
8. Repeat 5-8 until all the sample is transferred from the cup to the cylinder.
9. Add Dl water (at room temperature) to the cylinder to bring the level up to the 1000 mL mark.
10. Prepare a blank by adding 50 mL of sodium hexametaphosphate to a new clean sedimentation cylinder.
11. Add sufficient DI water to bring the level up to the 1000 mL mark.
12. Cover each cylinder with a watch glass and let stand overnight to equilibrate to room temperature (20°C to 25°C).
13. Before continuing the procedure, ensure that sufficient time is available to fully complete steps 14 to 25 (~8 hours for soils with high clay content).

*Note: Samples that have sat overnight and still have soil particles in suspension should be processed first*

14. Insert the plunger close to the bottom of the cylinder, and stir the suspension vigorously for about 2 minutes (approximately 5 strokes) by moving the plunger up and down the entire length of the column, taking care not to remove the plunger entirely from the suspension (otherwise bubbles will form, disrupting sedimentation). Also, use caution when nearing the top of the cylinder to avoid splashing and spilling the contents.

*Note: When mixing, ensure that no sediment remains in the lower 'corners' of the cylinder. In order to loosen and dislodge any sediment settled at the bottom of the cylinder, use short, strong, upward strokes, with the plunger near the bottom, and/or spin the plunger with the disk positioned just above the sediment.*

15. Once the suspension is stirred thoroughly, finish with two or three slow, smooth strokes and then remove the plunger, tipping it slightly to return adhering drops of suspension to the column.
16. Start timing as soon as stirring is finished (the settlement of particles begins immediately at the cessation of stirring) and do not wait until the plunger has been fully removed and cared for.
17. If the surface of the suspension is foaming, add 1-2 mL (drops) of amyl alcohol or apply a silicone spray.

*Note: For organic soils in addition to foaming there may be floating particles. 5.18 Immediately lower the hydrometer, very carefully, into the suspension, so that the hydrometer will not move up and down after releasing it.*

18. Exactly 40 seconds after stirring is finished, take the hydrometer reading at the top of the meniscus to the nearest 0.5 g.

*Note: Make sure that there are no air bubbles within the suspension. Air bubbles will affect the readings.*

19. Remove the hydrometer and insert the thermometer: read the temperature (to the nearest 0.5 °C) of the suspension.
20. Record the hydrometer and temperature readings
21. Clean the hydrometer and thermometer with DI water and wipe dry.
22. Repeat 10 to 20 for the control (Blank) cylinder.
23. Let the cylinders stand undisturbed.
24. At the end of 2 hours (120 min) take hydrometer and temperature readings.

*Note: For high clay soils repeat reading after 7 hours. If different from the2-hour reading use the 7-hour reading. Keep the 2-hour reading for the blank.*

25. The corrected 40 second measurement (M40) is the sum of the hydrometer reading (R40) minus the control reading (C40) at 40 seconds and the room temperature (RT) minus 20 multiplied by 0.36 (equation E1). The corrected 2-hour measurement (M2) is the sum of the hydrometer reading (R2) minus the 2-hour control reading (C2) and the room temperature minus 20 multiplied by 0.36 (equation E2). The clay content (% Clay) is calculated by dividing the initial sample weight (W) by the corrected 2-hour measurement multiplied by one hundred (equation E3). The slit and clay content (% Slit&Clay) is measured by dividing the initial sample weight (W) by the corrected 40 seconds measurement multiplied by one hundred (equation E4). The slit (%Slit) content is calculated by subtracting the value for the clay content from the slit and clay content (equation E5). Finally, the sand (% Sand) content is calculated by subtracting the value for the slit and clay content from 100 (equation E6).

Equation E1.

$$M40 = (R40 - C40) + [(RT - 20) * 0.36]$$

Equation E2.

$$M2 = (R2 - C2) + [(RT - 20) * 0.36]$$

Equation E3.

$$\%Clay =(W/M2)* 100$$

Equation E4.

$$\%Slit\&Clay = (W/M40)* 100$$

Equation E5.

$$\%Slit = \%Slit\&Clay - \%Clay$$

Equation E6.

$$\%Sand = 100 -\%Slit\&Clay$$
